# Virus-associated organosulfur metabolism in human and environmental systems

**DOI:** 10.1101/2021.01.05.425418

**Authors:** Kristopher Kieft, Adam M. Breister, Phil Huss, Alexandra M. Linz, Elizabeth Zanetakos, Zhichao Zhou, Janina Rahlff, Sarah P. Esser, Alexander J. Probst, Srivatsan Raman, Simon Roux, Karthik Anantharaman

## Abstract

Viruses influence the fate of nutrients and human health by killing microorganisms and altering metabolic processes. Organosulfur metabolism and biologically-derived hydrogen sulfide play dynamic roles in manifestation of diseases, infrastructure degradation, and essential biological processes. While microbial organosulfur metabolism is well-studied, the role of viruses in organosulfur metabolism is unknown. Here we report the discovery of 39 gene families involved in organosulfur metabolism encoded by 3,749 viruses from diverse ecosystems, including human microbiomes. The viruses infect organisms from all three domains of life. Six gene families encode for enzymes that degrade organosulfur compounds into sulfide, while others manipulate organosulfur compounds and may influence sulfide production. We show that viral metabolic genes encode key enzymatic domains, are translated into protein, are maintained after recombination, and that sulfide provides a fitness advantage to viruses. Our results reveal viruses as drivers of organosulfur metabolism with important implications for human and environmental health.

## Introduction

Biological sulfur cycling is one of the oldest and most influential biochemical processes on Earth and is primarily driven by microbial reduction of sulfate to produce hydrogen sulfide (Andreae, 1990; Fike et al., 2015; Wacey et al., 2011). Sulfide plays dynamic roles in the degradation of infrastructure and souring of oil reserves (Ma et al., 2000; Voordouw et al., 1996), microbial respiration and essential biosynthesis processes, and manifestation of human gastrointestinal disorders such as colitis, inflammatory bowel diseases (IBD) and colorectal cancer (CRC) (Guo et al., 2016). Much of our knowledge of sulfur cycling focuses on a small subset of microbes that are capable of respiring inorganic sulfur compounds, a process known as dissimilatory metabolism (Anantharaman et al., 2018). Consequently, the cycling of sulfur-containing organic (organosulfur) compounds and resulting sulfide production from more widespread biological mechanisms and sources has largely been ignored.

Two mechanisms of sulfide production include the degradation of organosulfur compounds and assimilatory sulfur metabolism. Sulfide production from microbial-driven degradation of organosulfur compounds, such as the amino acid cysteine, has been noted as a significant contributor to sulfide concentrations in environmental and human systems (Carbonero et al., 2012; Morra and Dick, 1991; Xia et al., 2017). However, there exists no comprehensive analysis of the specific microbes involved. Assimilatory sulfur metabolism, a common strategy used by many microbes and some eukaryotes to incorporate sulfide into biological compounds, has similarly been routinely discounted as a mechanism of significant sulfide release into either environmental or human systems. Notably, the role of viruses in these processes has not been explored.

Microbial viruses, mainly comprising bacteriophages (phages) are extraordinarily abundant on Earth. Microbial viruses are known to redirect and recycle nutrients on the scale of ecosystems by infecting and lysing host cells (Gobler et al., 1997; Jiao et al., 2010; Jover et al., 2014; Wilhelm and Suttle, 1999). In the oceans alone, the number of viral infections per second exceeds the number of stars in the known universe, which likely leads to the lysis of over 20% of all microbes per day (Manojlović, 2015; Suttle, 2007). In addition to lysis, viruses can actively redirect host metabolism during infection which manipulates major biogeochemical cycles, including carbon, nitrogen and sulfur. One such mechanism involves viruses “stealing” metabolic genes from their host in order to gain fitness advantages during infection (Sullivan et al., 2006). Such host-derived viral genes are termed auxiliary metabolic genes (AMGs), and are expressed during infection to modulate microbial respiration, biosynthesis processes, and/or direct intracellular nutrients towards virus replication and virion production (Anantharaman et al., 2014; Breitbart et al., 2007; Hurwitz et al., 2013, 2015; Mann et al., 2003; Roux et al., 2014; Suttle, 2005; Thompson et al., 2011; Trubl et al., 2018). For example, some viruses of Cyanobacteria encode core photosystem proteins that augment host metabolism in order to increase the biosynthesis of dNTPs that are utilized for viral genome replication (Thompson et al., 2011). The viral auxiliary metabolism of iron-sulfur clusters, central carbon metabolism, nitrification, methane oxidation and other metabolic processes could also provide viruses with a multi-faceted method of manipulating nutrients within their host cell to enable efficient, rapid or otherwise a more improved viral replication cycle (Ahlgren et al., 2019; Chen et al., 2020; Hurwitz and U’Ren, 2016; Hurwitz et al., 2015).

In spite of the importance and global prevalence of viruses, nothing is known about their contribution and impact on AMG-driven organosulfur metabolism in the environment. Moreover, the role of AMGs in human microbiomes has been largely unexplored. Here, we investigated environmental and human microbiomes for the presence of viruses involved in production of hydrogen sulfide and manipulation of organosulfur metabolism. By screening publicly available partial and complete viral genomes from cultivated and uncultivated viruses, we identified genes involved in direct and indirect sulfide production from organosulfur degradation and assimilatory sulfur metabolism. We followed this up with experiments to validate the impacts of genes for organosulfur metabolism as well as hydrogen sulfide on viral fitness.

## Results

### Metabolic pathways for organosulfur metabolism driven by viral AMGs

We queried a comprehensive dataset of approximately 135,000 partial and complete viral genomes (contigs) publicly available on Integrated Microbial Genomes/Viruses (IMG/VR) (Paez-Espino et al., 2016, 2017) and the National Center for Biotechnology Information (NCBI) databases, and two metagenomic studies from Lake Mendota, WI (Linz et al., 2018), for the presence of virally encoded proteins for organosulfur metabolism. In total, we identified 4,103 viral AMGs representative of 39 unique gene families. All genes identified are categorized as Class I AMGs, or those for central metabolic functions but auxiliary to productive viral infection (Hurwitz and U’Ren, 2016). These AMGs were detected on 3,749 non-redundant viral genomes from all major bacterial dsDNA viral families (*Myoviridae, Podoviridae* and *Siphoviridae*) including viruses infecting an archaea (Rahlff et al., 2020) and eukaryote (amoeba) (Schulz et al., 2020). Therefore, AMGs for organosulfur metabolism were identified on viruses infecting all three domains of life, representing a shared metabolic constraint regardless of host domain. The viruses represent cultivated and uncultivated viruses, linear and circular genomes, and lytic and lysogenic cycles of viral replication across a vast range of environmental and human microbiomes. Of these, 164 have been isolated and cultivated on hosts spanning nine major bacterial lineages (Alphaproteobacteria, Betaproteobacteria, Gammaproteobacteria, Cyanobacteria, Actinobacteria, Firmicutes, Bacteroidetes, Verrucomicrobia and Deinococcus-Thermus) as well as an amoeba (*Vermamoeba vermiformis*) (**Table S1**). The isolation of viruses encoding organosulfur metabolism AMGs indicates that the identification of such viral driven metabolism is not an artifact of metagenomic analysis.

Viral AMGs are putatively associated with five distinct processes: sulfide production from organic sulfur, the assimilatory sulfate reduction pathway, sulfite production from organic sulfur, metabolism of organic sulfur, and sulfur-related amino acid metabolism (**Figure 1 and Table 1**). Six different AMG families (*cysK, cysM, malY, dcyD, metC* and *metY*) encode for enzymes able to directly produce sulfide from the degradation of cysteine and homocysteine, which are important organosulfur compounds and central sources of sulfur in the environment and human body (Chiku et al., 2009; Fitzgerald, 1976). Six other AMG families (*cysD, cysN, cysC*, bifunctional-*cysNC*, *cysH* and *cysJ*) are components of the assimilatory sulfate reduction pathway, which is widely utilized across all three domains of life for incorporation of sulfide into cysteine. Sulfite can be directly produced from the breakdown of several organosulfur compounds (e.g. taurine) by three families of AMGs (*tauD, ssuD* and *msmÄ*) and successively fed into dissimilatory and assimilatory sulfate reduction. Eleven of the AMG families (*aspB, metB, metH, metE, msrC, metK, megL, dcm, mtnN, ahcY* and *luxS*) are inferred to indirectly produce sulfide by manipulating abundant organosulfur compounds (e.g. methionine and cystathionine) that funnel into the synthesis of cysteine or homocysteine. Finally, indirect organosulfur metabolism by the remaining thirteen AMG families (*lysC, thrA, asd, hom, metA, cysE, cysQ, nrnA, speE, mdh, mtnD, mtnA* and *mtnK*) would influence the synthesis of organosulfur compounds (e.g. synthesis of cysteine using serine) that feed into sulfide producing reactions.

**Figure 1.**
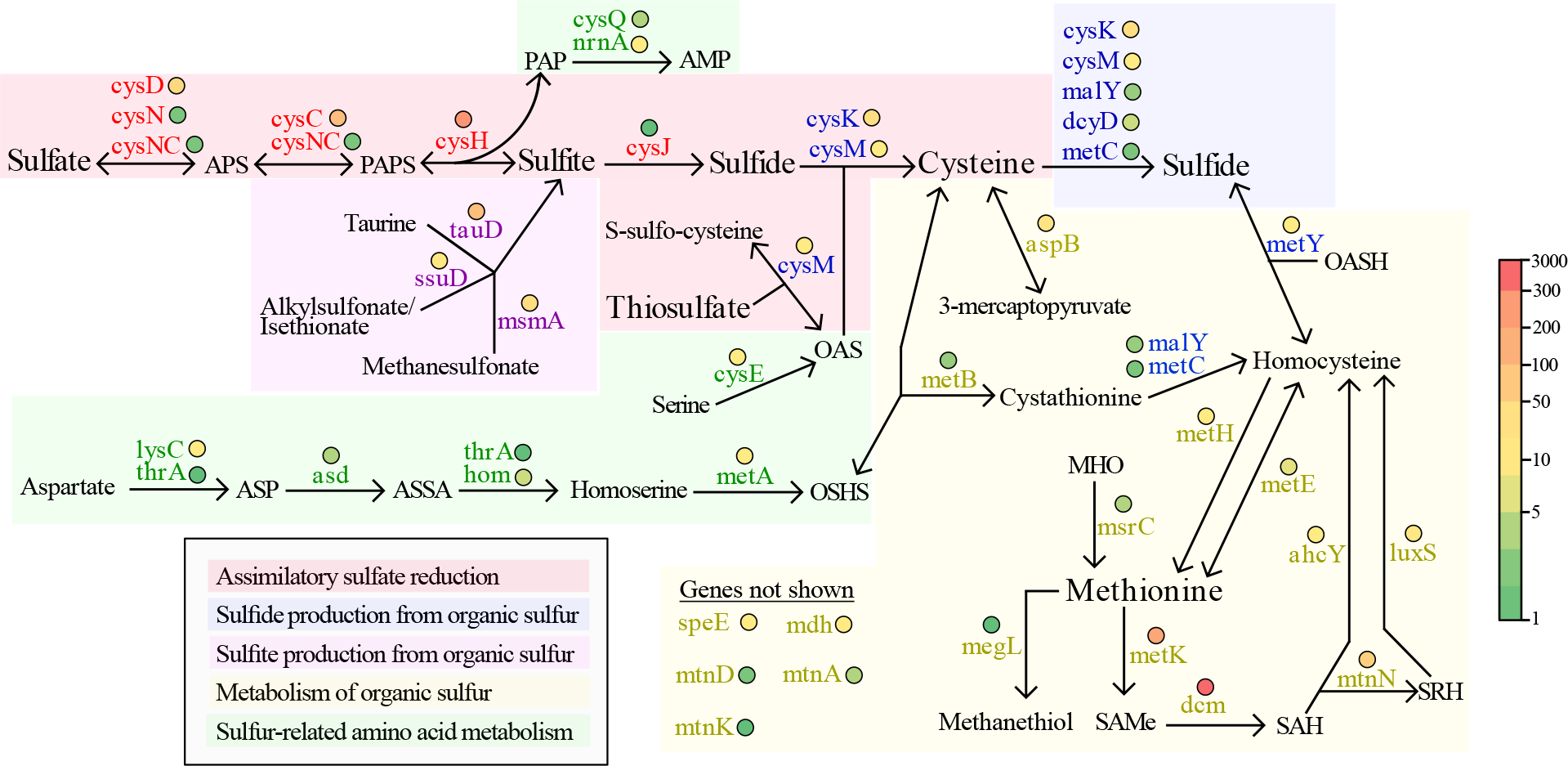
Reaction diagram of organosulfur transformations mediated by viruses. All genes shown have been identified on viruses and are colored coordinated respective to the process with which they are putatively associated. Colored circles represent the abundance of each AMG across all viral genomes according to the color scale (heatmap) on the right. Complete reactions and full names of acronyms are listed in Table 1.

**Table 1.**
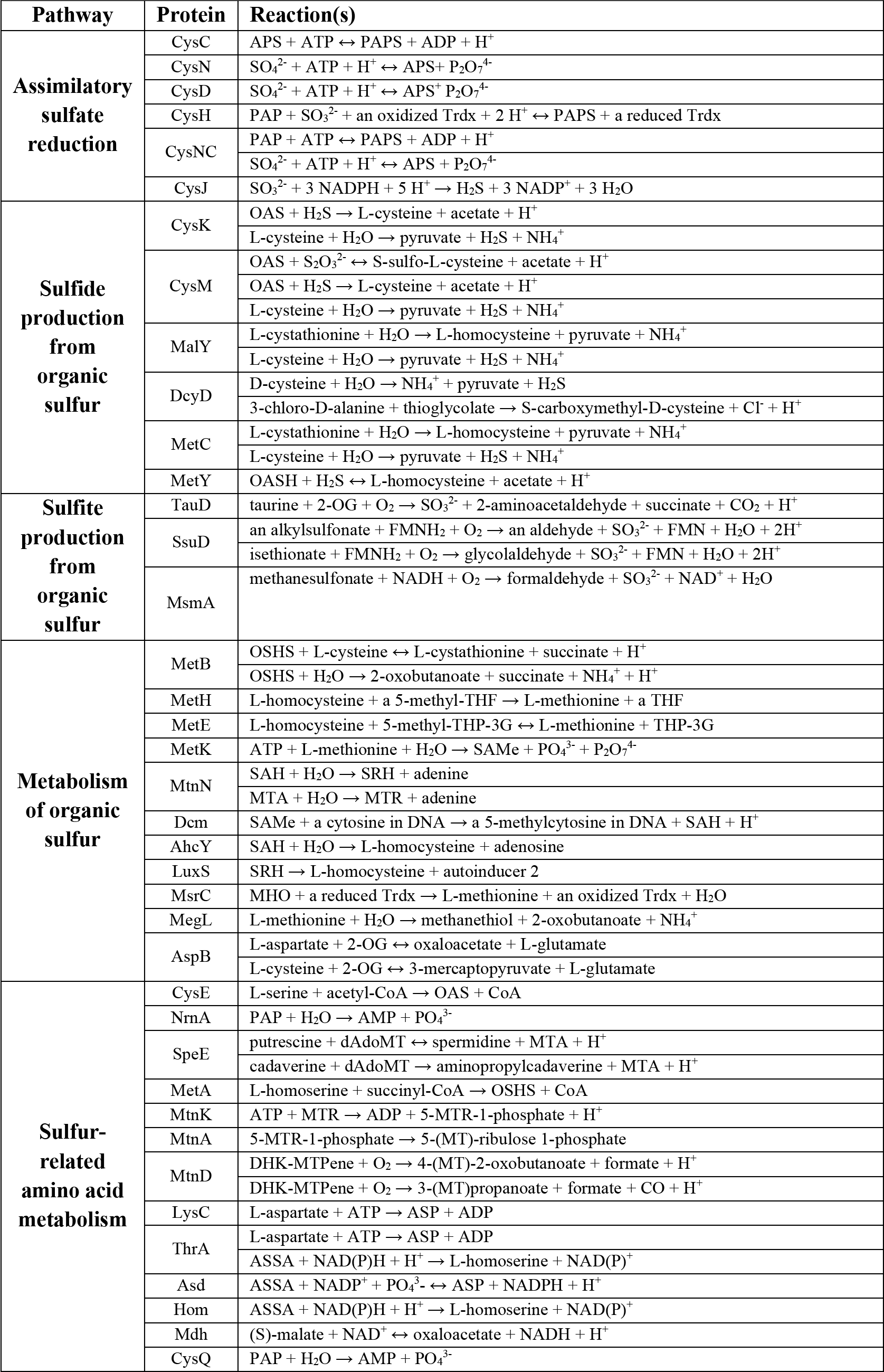
Complete reaction(s) performed by each AMG-encoded protein. Each protein is grouped respective to the main organosulfur metabolism pathway in which it is involved. Full names of acronyms are as follows. PAP: adenosine 3’,5’-bisphosphate, APS: adenosine 5’-phosphosulfate, PAPS: 3’-Phosphoadenosine-5’-phosphosulfate, CoA: Coenzyme A, OG: oxoglutarate, OAS: O-acetyl-L-serine, OASH: O-acetyl-L-homoserine, OSHS: O-succinyl-L-homoserine, SAMe: S-adenosyl-L-methionine, dAdoMT: S-adenosyl 3-(methylsulfanyl)propylamine, MTA: S-methyl-5’-thioadenosine, MTR: 5-(methylsulfanyl)-α-D-ribose, MT: methylsulfanyl, SAH: S-adenosyl-L-homocysteine, SRH: S-ribosyl-L-homocysteine, DHK-MTPene: 1,2-dihydroxy-5-(methylsulfanyl)pent-1-en-3-one, ASSA: L-aspartate 4-semialdehyde, ASP: L-aspartyl-4-phosphate, MHO: L-methionine-(R)-S-oxide, Trdx: thioredoxin, THF: tetrahydrofolate, THP-3G: tetrahydropteroyl tri-L-glutamate.

### Viruses encoding AMGs for organosulfur metabolism are globally distributed

Uncultivated viruses encoding AMGs for organosulfur metabolism were recovered from diverse environmental (marine, freshwater, engineered, soil, hydrothermal vent, non-marine saline and alkaline, deep subsurface, wetland and thermal spring), non-human host-associated (mammalian gut, other animal-associated and plant-associated) and human host-associated (gastrointestinal, oral and vaginal) microbiomes (**Figure 2A**). Cultivated and well-characterized viruses exhibited likewise microbiome dispersal because they were recovered from more than one ecosystem (e.g. food production, marine, freshwater, soil, engineered, hot springs, animal-associated, plant-associated, as well as human-associated gastrointestinal, oral and skin) (**Table S1**). These results encompassed every ecosystem category, with the exception of air, in which viruses are routinely identified. This displays evidence that viruses encoding AMGs for sulfide production are ubiquitous on Earth.

**Figure 2.**
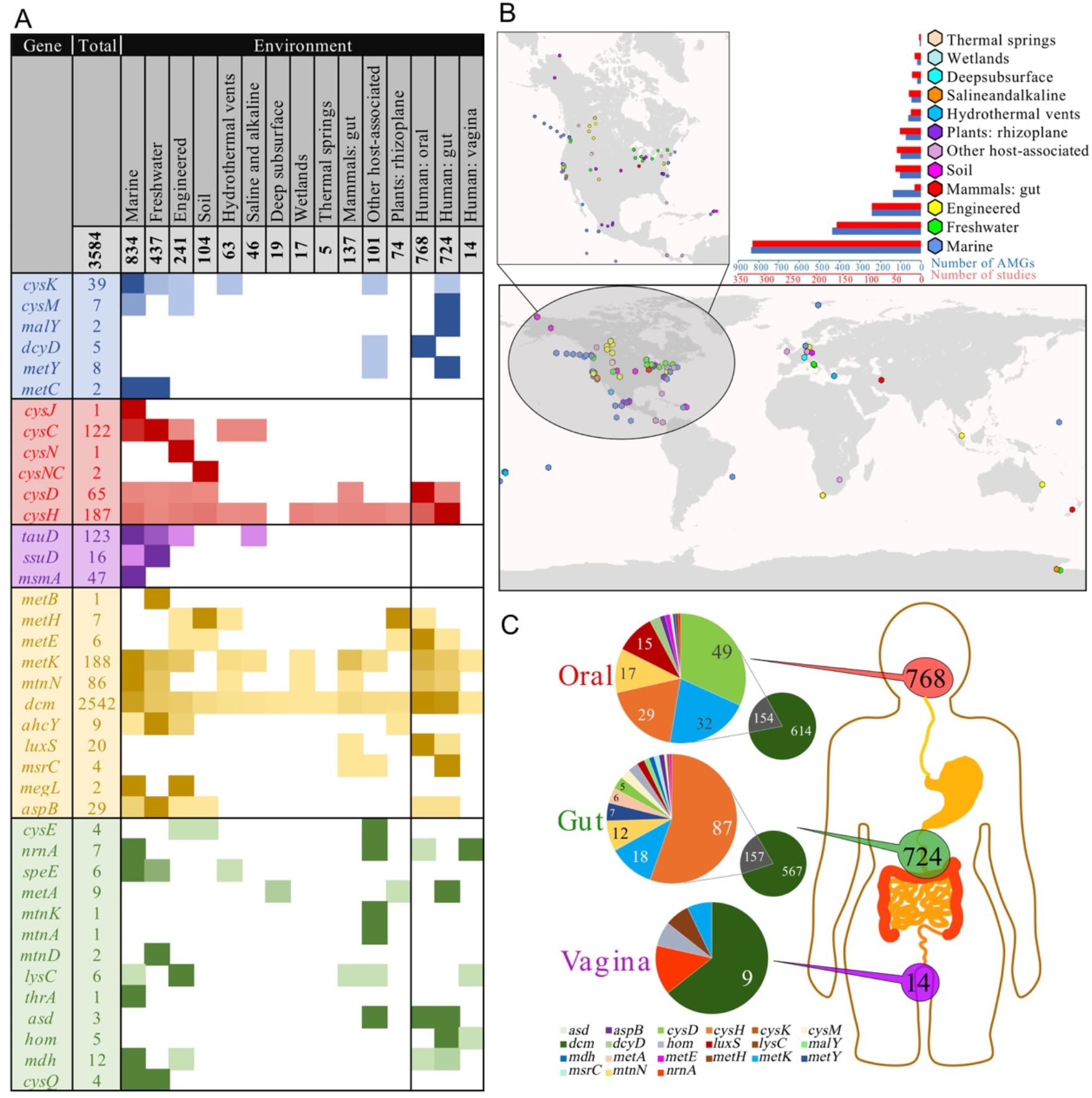
Distribution of viral AMGs in environmental and human microbiomes. (A) Heatmap of each AMG’s relative abundance in environmental and human systems with colors coordinated by the AMG’s pathway respective to Figure 1. Per AMG, darker colors represent greater abundance. A total of 3,584 AMGs derived from IMG/VR are shown. (B) Global distribution of viruses encoding AMGs, color coordinated by environment classification. The bar graphs represent the number of AMGs and IMG studies from which viruses were recovered. See Tables S2 and S3 for exact abundances for (A) and (B), respectively. Only studies with published coordinates and environment categories are shown. (C) Abundance of AMGs derived from incomplete or uncultivated viruses from human oral, gastrointestinal and vaginal microbiomes. Only values greater than five are shown.

Next, we estimated the proportion of viral richness in each ecosystem category found to encode organosulfur metabolism AMGs. Viruses encoding at least one AMG were found to be highly abundant in human vaginal, gastrointestinal and oral microbiomes comprising 8%, 6% and 3% of all identified viruses, respectively. Mammalian-associated, other animal-associated and plant-associated microbiomes likewise had significant AMG-encoding virus abundances of 8%, 6% and 6%, respectively. Notably, previous reports have determined that expanded viral richness in the gastrointestinal tract is correlated with the manifestation of IBD (Norman et al., 2015) and our results support the possibility of this being in part due to the metabolic potential of viruses, such as for sulfide production. This points to an important distinction that the collective metabolic potential of viruses in these host-associated environments, in conjuction with measuring total viral richness, could have significant implications for host health. Viruses encoding organosulfur AMGs beyond host-associated microbiomes may also impact ecosystem health. Major environmental systems, such as the deep subsurface (6%), engineered (3%), soil (3%), freshwater (2%), wetlands (2%), marine (2%) and hydrothermal vents (2%), likewise display significant richness of organosulfur AMG encoding viruses (**Table S4**). The net impact of viral metabolism on organic and inorganic sulfur compound concentrations in these environments is unknown, but it is nonetheless striking that up to 8% of all resident viruses may be involved.

Viruses recovered from non-human microbiomes also displayed extensive geographical and niche distributions, which demonstrates their relevance in global sulfur biochemistry (**Figure 2B**). Individual distributions of abundant AMGs (e.g. *dcm*, *cysC, cysK, cysH, metK*, and *tauD*) likewise had no geographical or environmental restrictions (**Figure S1A-F**). For example, *cysH* which encodes a critical enzyme for assimilatory sulfur metabolism was found in every ecosystem except the deep subsurface. CysK, a predominant enzyme involved in sulfide generation from cysteine degradation was also broadly dispersed in marine, freshwater, engineered, hydrothermal vent and host-associated environments. Even *msmA* which was only identified in marine environments showed strong geographical dispersal (**Figure S1G**).

AMG distributions between environments may depend on different factors, such as how universal the AMG function is (e.g. CysH and CysK are common amongst bacteria) or the nutrient landscape in a specific environment (e.g. MsmA is capable of degrading methanesulfonate, a common compound in marine environments (Henriques and Marco, 2015). However, human-associated samples contained the greatest fraction of identified *cysH* and *cysD* AMGs overall, while marine and freshwater environments contained nearly all of the identified *cysC*. In human-associated samples, nearly 97% of AMGs were *cysD, cysH, metK, mtnN, luxS* and *dcm* which encompass essential steps of cysteine and methionine degradation (**Figure 2C**). The uneven distribution of these assimilatory sulfate reduction AMGs suggests that further constraints on nutrient availability or variance in rate limiting steps based on thermodynamics in different environments play a role in determining the distribution of organosulfur metabolism AMGs.

### Viral organosulfur AMGs result in likely functional proteins and provide a fitness advantage to the virus

To overcome the challenge of assigning conclusive function to protein sequences in the absence of biochemical evidence, we analyzed functional and conserved domains of AMG-encoded proteins with biochemically characterized bacterial homologs. Overall, we examined 24 AMG families and found broad conservation of whole protein sequence and functional amino acid residues (**Figure S2**). For example, viral sequences encode specific domains for: CysC: ATP binding (gsGKss) and required motifs (dgD) (Poyraz et al., 2015); CysK: cofactor pyridoxal phosphate binding (KDR, NtG, GT/SgGT and SS/AG), substrate binding (T/SSGN and QF) and phosphate recognition (GI/V) (Ishikawa et al., 2010); MetK: substrate binding (egHPDk, acE, gEit, GDqG, DaK, TgRKi, sGKd and kvDrs) (Komoto et al., 2004); CysH: iron-sulfur cluster motif (CC…CxxC) (Chartron et al., 2006); TauD: nitrogen and oxygen binding (e.g. WH and H) (Knauer et al., 2012). Conserved amino acid residues that are not functional are likely preserved for structural features. The retention of AMGs on viral genomes despite strong selective pressures for reduced genome size suggests that most of these AMGs are functional (Bragg and Chisholm, 2008). In addition to functional and conserved domain analysis we calculated the ratio of non-synonymous to synonymous nucleotide differences (*dN/dS*) for a subset of the abundant viral AMG families. A *dN/dS* value less than one would suggest that the virus is under selective pressures to retain a functional AMG. *dN/dS* calculations for *cysK, cysC, cysD, cysH, tauD, msmA, metK, mtmN and luxS* AMG pairs revealed that viral AMGs appear to be under purifying selective pressures to retain function of the encoded AMGs (Supplementary Figure S3).

To assess if viral AMGs are active in the environment, we queried a comprehensive metagenomic and metatranscriptomic dataset from Lake Mendota, WI. We identified 23 AMGs representative of six gene families (*aspB*, *cysC*, *cysH*, *metK*, *speE* and *tauD*) that were actively expressed by 22 different viruses over a 48-hour time period (**Table S5**). One *cysC* in particular was expressed by a virus with a 210kb genome that was bioinformatically determined to be complete and circular. Analysis of the genome’s GC-skew, a metric to evaluate genome replication patterns using nucleotide coverage (Sernova and Gelfand, 2008), was used to determine that the virus performs rolling circle replication (i.e. unidirectional) which is a common method utilized by viruses (Olm et al., 2017a) (**Figure S4A**). To assess if the virus was actively replicating when *cysC* was expressed we used a metagenomic read mapping approach to estimate the genome’s *in situ* index of replication (iRep) (Brown et al., 2016). The genome’s iRep value of 1.54 falls within the range of typical values of growing populations and indicates that the virus was actively replicating its genome in the environment when *cysC* was expressed (**Figure S4B**). Analyses of other host-virus systems with transcriptomic data enabled the identification of *cysH* expression by Enterobacteria phage Lambda during infection of *Escherichia coli* MG1655 (Liu et al., 2013). The activity and expression of viral AMGs in various systems provides further evidence that they are likely utilized for a specific function during infection.

To validate that AMGs are in fact transcribed during infection we developed a model host-virus system with *Lactococcus lactis* C10 and its *cysK*-encoding virus Lactococcus phage P087. The transcript abundance of *cysK* was measured in a culture of either *L. lactis* C10 grown alone (control) or with P087 at timepoints 15-, 60- and 120-minutes post infection (**Figure S5** and **Table S6**). At 120 minutes the host cells in the infection condition had mostly lysed from viral infection. Transcript abundance of *L. lactis* C10 *cysK* was found to be comparable at 15 minutes and 60 minutes in either the uninfected control or infected with P087. At 120 minutes transcripts of *L. lactis* C10 *cysK* were 4x greater than at 60 minutes in the control but were undetectable in the infected condition. This suggests that *L. lactis* C10 *cysK* transcripts are greatly reduced during mid to late infection by P087. The transcript abundance of P087 *cysK* follows a similar trend as *L. lactis* C10 *cysK*. At 15 minutes P087 *cysK* transcripts were near zero and by 60 minutes were in approximately 2x greater abundance compared to transcripts of the host. By 120 minutes P087 *cysK* transcripts likewise reduced nearly to initial levels. There was no detection of P087 *cysK* transcripts within the uninfected control. Although P087 *cysK* transcript abundance never exceeded that of *L. lactis* C10 *cysK*, we provide further evidence that the viral AMG *cysK* is actively transcribed during infection and potentially replaced host *cysK* to an extent with the greatest abundance during mid infection rather than early or late infection.

To validate that transcribed AMGs in fact produce protein, we further leveraged the *L. lactis* and P087 system. Using untargeted mass spectrometry at the endpoint of virus infection (i.e. lysis) we identified that P087’s AMG *cysK* produces protein and at approximately 1.5x greater abundance than *L. lactis* C10 *cysK* (**Table S7**). The higher ratio of virus CysK to host CysK suggests the virus gains a fitness advantage from compensation of CysK levels in the cell. These findings build upon the results from our qPCR based analysis of transcript abundance in which host transcripts were more abundant than viral but may be explained by higher stability of either viral CysK or *cysK* transcripts. Moreover, since viruses demand a substantial fraction of cellular resources during infection (Mahmoudabadi et al., 2017), the high viral CysK levels measured here supports our hypothesis that CysK is actively utilized during productive infection in contrast to being metabolically inactive. The presence of the gene on the genome in conjunction with transcription and translation measurements is consistent with the AMG providing a fitness advantage, which has been modeled to be as much as a 4% gain for some AMGs (Bragg and Chisholm, 2008). The mechanism(s) by which this functions is likely different than what has been observed previously for AMGs. For example, AMGs for photosynthesis were found to have differential effects during light-dark cycles as well as transcript compensatory effects over an ~8 hour time period (Thompson et al., 2011). Conversely, P087 is not influenced by light-dark cycles and complete lysis can occur within ~2.5 hours. Beyond providing evidence that AMGs can be remarkably active during infection this further underlines the diverse nature by which AMGs are utilized by viruses. In addition, the identification of similar gene families on genomes of diverse, geographically spread viruses strongly supports the hypothesis that organosulfur metabolism AMGs play a functional role during infection (Roux et al., 2014).

### Viruses encoding organosulfur AMGs are phylogenetically diverse

To investigate the diversity of AMGs we conducted phylogenetic analysis of encoded amino acid sequences for five gene families. Phylogeny of CysH from complete viral genomes show close relationships between viruses and their known hosts, supporting previous observations that AMGs are most often acquired from the host (Sullivan et al., 2006) (**Figure S6A**). One clade in particular encoded an addition domain of unknown function (DUF3440) which suggests a shared evolutionary history. Analysis of CysH phylogeny of viral contigs with no known host revealed a similar clustering of viruses with their putative bacterial hosts (phyla Bacteroidetes and Firmicutes) (**Figure S6B**). In contrast to CysH, phylogenetic analysis for several abundant AMG protein sequences (CysC, CysK, TauD and MetK) on complete and incomplete viral genomes displayed clustering of viral sequences in separate clades from bacterial homologs with few exceptions of the virus clustering with a putative host (**Figure S6C-F**).

Separate clustering would suggest that viruses may have acquired AMGs beyond their current or known host range, which is supported by the observation that viruses can encode an AMG that their host does not (e.g. *cysC* for *Xylella* phage Sano) and that AMGs can cluster separately from their host (e.g. CysH for *Vibrio* phages). However, based on the CysH phylogeny of complete viral genomes another likely explanation for distinct viral clustering is that the full range of host sequences has yet to be identified. Within the human microbiome alone, thousands of novel bacterial genomes have been identified recently and may provide further insight into host ranges or origins of AMG transfer (Almeida et al., 2019; Nayfach et al., 2019; Pasolli et al., 2019). Even so, in comparison to human microbiomes, little is known about the breadth and diversity of environmental or human viromes. Analysis of all AMGs suggests they have collectively been derived from bacteria (with the exceptions of the archaeal and eukaryotic virus) affiliated with the phyla Firmicutes, Bacteroidetes, Alphaproteobacteria and Gammaproteobacteria, which is supported by the host range of cultivated AMG-encoding viruses (**Figure S7**).

### Directed recombination and AMG sequence conservation validates proposed mechanism of AMG transfer and retention

The proposed mechanism of AMG acquisition by viruses in nature is the transfer of a host metabolic gene to the virus by recombination. Over multiple replication cycles of the viral genome, the AMG is retained as a functional gene. To verify this proposed mechanism, we engineered *Escherichia coli* phage T7 by inserting the host gene *cysK* (T7::*cysK*) to simulate a recombination event. Following successful insertion, T7::*cysK* was passaged, in three biological replicates, for nine complete infection cycles to simulate infection in nature over time. After passaging, the T7::*cysK* construct was sequenced to check for retention of the AMG in the viral population. Sequencing confirmed retention of the gene, indicating that recombination of a host metabolic gene onto a viral genome (i.e., AMG acquisition) can lead to stable retention of an AMG over time. Furthermore, between three biological replicates no mutations from the wildtype *cysK* sequence were observed.

Importantly, these observations show that a recombination event can occur without environmental triggers (e.g., nutrient limitation during infection) or fitness constraints (e.g., metabolic bottlenecks in the host), which provides further credibility for the proposed mechanism that AMG transfer occurs frequently and randomly in nature. If the AMG provides sufficient fitness benefits, or a lack of detrimental effects on viral replication it will be retained over multiple infection cycles. In the system developed here, conditions resulting in a fitness benefit (e.g., greater burst size or faster replication) for the T7::*cysK* virus compared to wild-type T7 were not identified.

### Sulfide can provide a fitness advantage to viruses

Since active expression and function of AMGs likely can result in the production of sulfide in the environment and human microbiome, we sought to determine if sulfide does indeed confer a fitness advantage to viruses. A highly plausible method for viruses to achieve this would be through the degradation of cysteine which is present in nearly all environments. As a result, we hypothesized the *cysK*-encoding virus P087 would have the capacity to gain a fitness advantage in the presence of sulfide. Theoretically P087 would be involved in the direct degradation of intracellular cysteine via the action of virally encoded CysK under some conditions. To elucidate if sulfide alone confers a fitness advantage, we exogenously added sulfide during P087 infection of *L. lactis* and quantified the impact on virus and host growth. We found that viable virus production increased linearly with the addition of physiologically relevant concentrations of sulfide (**Figure 3A**) with no significant observed differences in host growth (**Figure 3B**). This indicates that under the conditions tested P087 benefits from increased production of sulfide in the system through either AMG or host-driven mechanisms, and that the resulting fitness gain is not due to a simple increase in host abundance. We performed the same experiment with exogenously added cysteine but did not observe any effect on viral fitness (data not shown). This has significant biological implications as microorganisms contain high intracellular concentrations of cysteine, with *L. lactis* species reported to contain approximately 3.5mM intracellular cysteine (Li et al., 2005). Likewise, *Escherichia coli* has a free cysteine pool of approximately 150μM (Park and Imlay, 2003). We believe other viruses encoding organosulfur metabolism AMGs would likewise derive a fitness advantage under similar conditions and that this phenotype is not restricted to the ability to directly produce sulfide from cysteine degradation.

**Figure 3.**
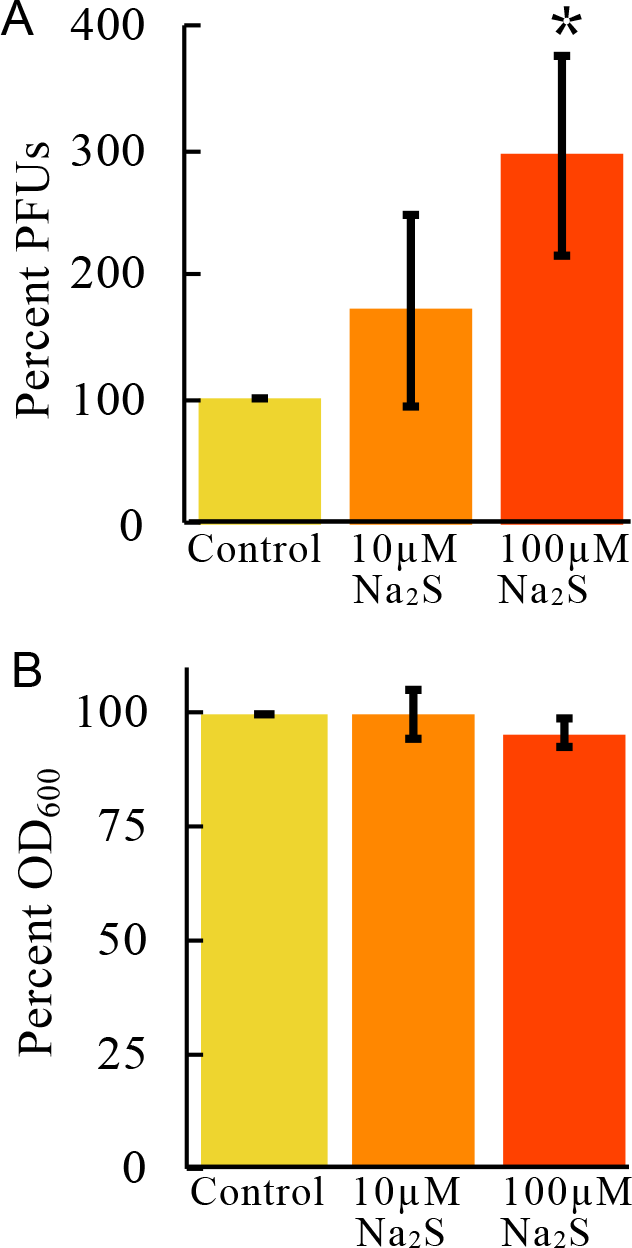
Increased viral fitness is associated with sulfide concentrations. Impact of varying sulfide concentrations on (A) Lactococcus phage P087 virus production as measured by plaque forming units (PFUs) and (B) uninfected host growth. Experimental conditions are normalized to percent of control. Asterisk represents statistical significance (p<0.02) compared to the control.

### Viral organosulfur auxiliary metabolism associated with human gut bacteria

Among viruses with known hosts, 107 were found to be associated with 35 different bacterial species known to be commensal or pathogenic residents of the human gastrointestinal tract (**Table S1**). These viruses encode five AMGs (*cysE, cysH, cysK, dcm* and *metK*) for both the assimilation of sulfur and capacity to degrade organosulfur compounds into sulfide. Most of these viruses were isolated from a variety of dairy, soil, sewage and wastewater environments indicating a potential for environmental reservoirs of sulfide producing viruses, or in the case of wastewater environments the viruses may have been resident in human gastrointestinal tracts. Five AMG-encoding viruses of the pathogens *Salmonella enterica, Staphylococcus aureus, Vibrio cholerae* and *Clostridium difficile* were isolated from human fecal samples indicating transmission and replication in human gastrointestinal tracts likely does occur and may contribute to dysbiosis via the production of sulfide or altering the organosulfur metabolic potential of the pathogenic host.

Uncultivated viruses from the human gastrointestinal tract encoding AMGs p utatively involved in direct sulfide production (*cysM, malY* and *metY*) had high protein identity (>97%) to *Alistipes putredinis, Alistipes obesi, Alistipes finegoldii, Bacteroides uniformis* and *Bacteroides vulgatus* suggesting they are viruses closely associated with these human gut bacteria from the order *Bacteroidales* (phylum Bacteroidetes) (Fenner et al., 2007; Hugon et al., 2013; Patrascu et al., 2017; Schirmer et al., 2018). Viruses encoding *metK, mtnN* and *metE* (i.e. capacity for methionine degradation to sulfide) in human gastrointestinal samples were likewise inferred to be closely associated with the human gut bacteria *Alistipes ihumii, Faecalibacterium prausnitzii, Flavonifractor sp., Bacteroides intestinalis, Bacteroides xylanisolvens, Bacteroides uniformis, Bacteroides thetaiotaomicron, Haemophilus parainfluenzae, Aggregatibacter sp*. and *Eubacterium sp*. based on high protein identity (Bakir et al., 2006; Costea et al., 2017; Curtis et al., 2014; Jiang et al., 2015; Kuang et al., 2017; Martín et al., 2017; Pfleiderer et al., 2014; Qin et al., 2010; Veiga et al., 2014). At lower protein identity (96%-80%), viruses encoding *metK, luxS* and *mtnN* were inferred to be in some part associated with the gut bacteria *Prevotella spp. (Bacteroidales), Butyricicoccus spp*. and *Clostridiales sp*. (Eeckhaut et al., 2013; Larsen, 2017; Patrascu et al., 2017) (**Table S8**).

Many of these *Bacteroidales* (i.e. *Alistipes spp., Bacteroides spp*. and *Prevotella spp*.) and some members of the phylum Firmicutes (e.g. *Haemophilus parainfluenzae* and *Butyricicoccus spp*.) have been strongly associated with IBD (Eeckhaut et al., 2013; Lucke, 2006; Schirmer et al., 2018; Veiga et al., 2014) and their role in inflammation may be in part attributed to virus-mediated or influenced production of sulfide. Importantly, viruses of these *Bacteroidales*, including *Prevotella* megaphages with high coding capacity, have been shown to be dominant and abundant in human gastrointestinal tracts which could promote the continuous viral-driven production of sulfide to exacerbate inflammation (Devoto et al., 2019; Dutilh et al., 2014).

### Comparative genomics displays diversity of viral genome organization

We used comparative genomics to examine the diversity of viruses found to be associated with human microbiomes. We identified four distinct uncultivated virus contigs encoding *dcm* from human oral samples to be closely related to known *Streptococcus pneumoniae* viruses based on genome sequence identity (**Figure S8A**). However, there are large stretches of dissimilarity between some of the genomes which may indicate evidence for large genetic exchange between viruses that frequently share the same niche and not the same host, which has been demonstrated before between *Lactococcus* and *Enterococcus* viruses (Villion et al., 2009). This observation supports the likelihood of AMG transfer between viruses in human and environmental microbiomes. Furthermore, two plant-associated viruses were identified to be closely related to known *Salmonella enterica* viruses originally derived from human fecal samples (**Figure 4A**). These plant-associated viruses may represent examples of environmental reservoirs for AMG-encoding viruses in the human gastrointestinal tract.

**Figure 4.**
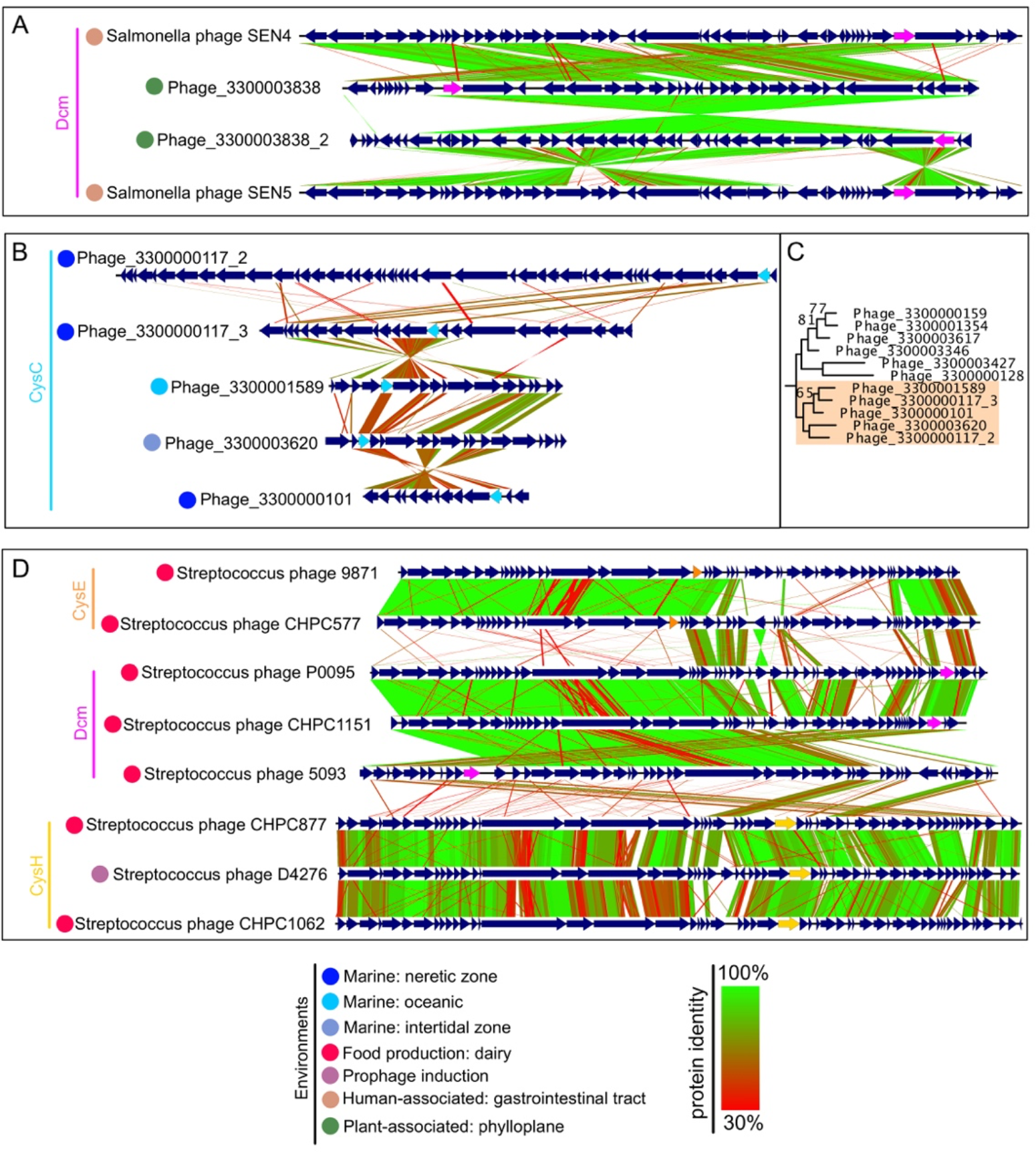
Genome comparisons of viruses encoding AMGs. Comparisons of (A) uncultivated viruses and complete *Salmonella enterica* viruses encoding *dcm* (pink), (B) uncultivated viruses encoding *cysC* (cyan) with (C) respective protein phylogeny (orange highlighting, refer to Figure S6 for full phylogenetic tree), and (D) complete *Streptococcus thermophilus* viruses encoding *cysE* (orange), *dcm* (pink) or *cysH* (yellow). For all comparisons, predicted open readings frames are annotated by dark blue arrows and genomes are connected with lines according to protein identity by tblastx alignment. Colored circles refer to the environment in which the virus was isolated or identified.

However, for either case above the exact nature of viral transfer of AMGs is challenging to determine because AMG sequences that closely share evolutionary history can be encoded on dissimilar and geographically diverse viruses. For example, five *cysC*-encoding viruses that group closely by CysC phylogeny conversely depict dissimilarity by genome comparison and are geographically dispersed in marine environments (**Figure 4B, C**). The same is true for six different *metK-*encoding viruses in which MetK shows phylogenetic similarity but the genomes are diverse and geographically spread (**Figure S8B**).

To further investigate the relationships of AMGs on viral genomes we examined the prevalence of multiple AMG copies on individual genomes. In total we identified 285 viral genomes that contained multiple copies. While most such genes encoded for identical functions (i.e. two copies of protein from the same gene family), some with connected (e.g. *metK* and *dcm*, *luxS* and *mtnN*) or disparate functions (e.g. *dcm* and *cysM, cysH* and *mtnN*) were also found. These findings suggest viruses may utilize these genes for diverse regulation of host organosulfur metabolism to fit their individual requirements (**Table S9**). For example, a single virus may augment both assimilatory sulfate reduction (e.g. using CysH) as well as methionine degradation (e.g. using MetK) during infection by encoding and expressing both AMGs.

We next compared viral genome organization to identify relationships in the physical location of AMGs between different viral genomes and interpret affiliations with other encoded genes. We found no universal organization of AMGs which were broadly encoded in various locations, such as between structural genes, adjacent to lysis factors, near genes for genome replication or nucleotide metabolism and within regions comprising genes of unknown function (**Figure 5**). Additionally, no pattern associated with encoding specific AMGs was detected according to virus classification, genome length or isolation source. There were a small number of outliers, such as a comparison of 10 complete viral genomes encoding *cysH* that indicated a trend towards co-location of the AMG with genome replication and/or nucleotide metabolism genes to suggest similar transcriptional regulation or function of this AMG across different viruses (**Figure S9**).

**Figure 5.**
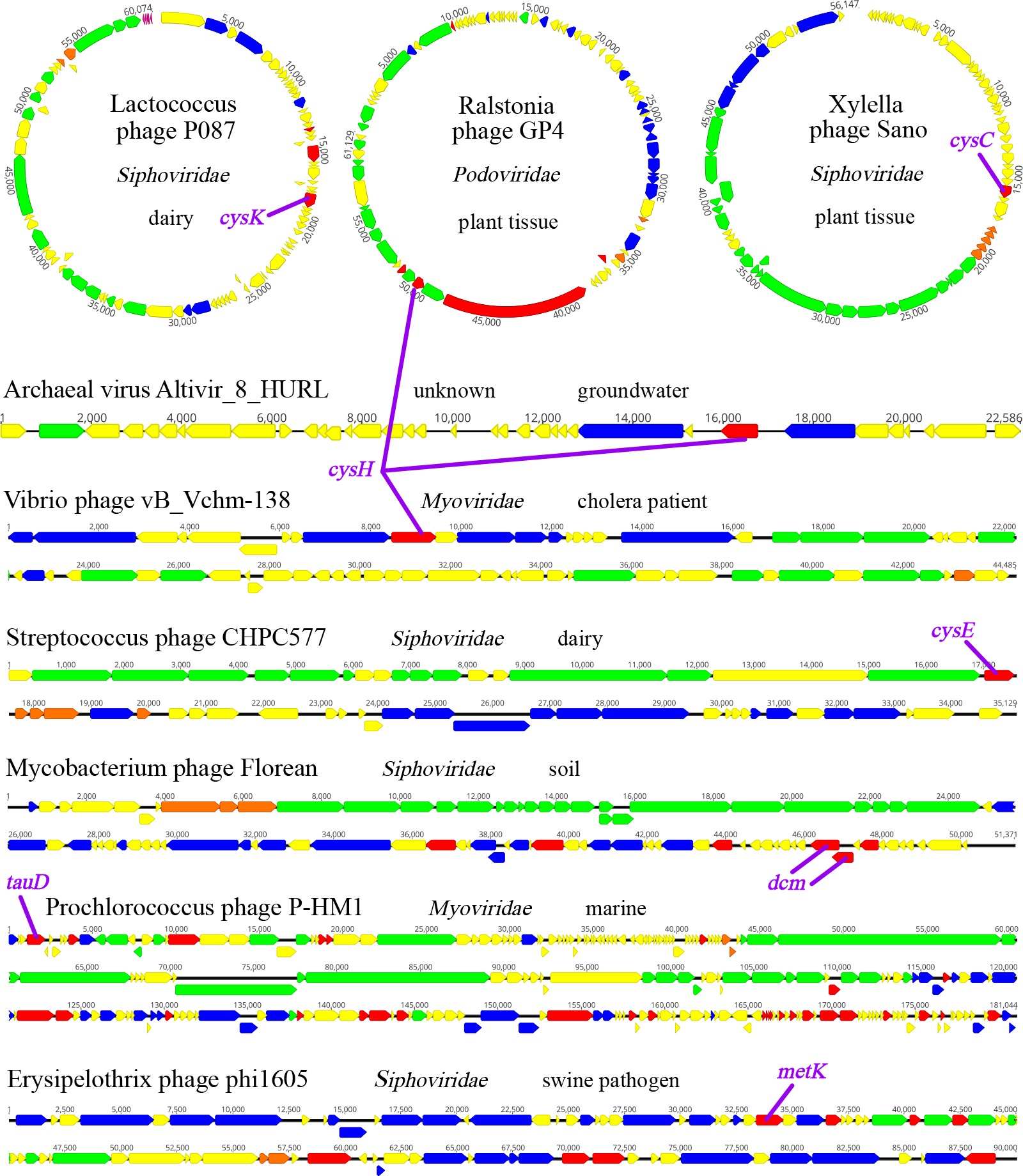
Genome organization of 8 complete viral genomes encoding organosulfur AMGs. Genome representation of circular and linear viruses. Arrows indicate open reading frames and are annotated by general function: virion structural assembly (green), auxiliary metabolism and general functions (red), nucleotide metabolism and genome replication (blue), lysis (orange) and unknown function (yellow). AMGs are annotated in purple.

The model that viruses acquire AMGs from diverse sources and for disparate functions is further supported by looking at AMG-encoding viruses that share the same host but not the same AMG. There are several different variations in which this occurs. One example involves *Bacillus cereus* phages PBC5, Basilisk, BCU4 and PBC6 where the viruses have low sequence similarity between genomes and AMG sequences (i.e. *cysH*) (**Figure S8C**). Another example involves *Streptococcus suis* phages phiJH1301-2, phiSC070807, phiNJ3 and phiD12 where the viruses have very similar genome sequences but encode multiple AMGs with similarity shared only among a subset of them (i.e. *metK* and *dcm*) (**Figure S8D**). A final example involves *Streptococcus thermophilus* phages 9871, CHPC577, P0095, CHPC1151, 5093, CHPC877, D4276 and CHPC1062 where the viruses group separately according to the single AMG each encodes (*cysE*, *cysH* or *dcm*) (**Figure S8D**). Taken together, these three examples indicate that viruses are able to employ separate strategies to accomplish a similar function of manipulating host organosulfur metabolism. This may be in the form of acquiring the same AMG from different sources to perform a shared task or acquiring disparate AMGs to perform separate tasks towards the same objective, such as sulfide production.

## Discussion

The metabolic potential of viruses, the most abundant biological entities on Earth, is all too often overlooked because viruses do not independently conduct metabolic transformations. Here we show that viral manipulation of host metabolism in contrast to solely measurements of viral richness and host range is likely important to the environmental sulfur cycle and human health. Furthermore, we propose that assimilatory sulfur metabolism, a ubiquitous method of fixing sulfur and manipulating organosulfur compounds, is frequently modulated by viruses during infection of organisms from all three domains, and in almost all microbiomes on Earth. This poses an important question, what have we been overlooking in viromes by frequently assessing sequence reads instead of metagenomically assembled genomes that encode AMGs? Are we giving enough emphasis on viruses as core drivers in the metabolism of microbiomes?

AMG-driven organosulfur metabolism mediated by viruses may lead to sulfide production in the gastrointestinal tract during infection or following microbial lysis. The result would be a sulfide-induced inflammatory response in conjunction with the activity of resident microbiota or invading pathogens, though the extent to which this occurs in human or environmental systems has yet to be quantified. Indeed, it has been observed that infected bacterial cells have manipulated and ‘rewired’ sulfur assimilation that will impact cysteine metabolism and likely sulfide production (Howard-Varona et al., 2020). Furthermore, viruses encoding sulfur assimilation AMGs may be short-circuiting the assimilatory sulfur pathway by reducing the steps necessary for assimilation of sulfur into organosulfur compounds. This concept is supported by the observation that *cysH* is the most abundant organosulfur metabolism AMG, which plays a role in both the canonical sulfate assimilation pathway as well as direct sulfonation of organic molecules (Moran and Durham, 2019). The latter mechanism may explain the high abundance of *cysH* on viral genomes.

The evidence presented here strongly points towards sulfide production as a component of viral organosulfur auxiliary metabolism, either directly or indirectly by AMG activity, which could provide many fitness advantages for viruses (**Figure 6A**). As obligate intracellular pathogens, viruses could benefit from the survival and enhanced growth of their host, which could be achieved by responding to sulfur starvation signals, assimilating sulfide for biosynthesis (e.g. for sulfolipids), upregulating sulfide utilization (e.g. sulfide oxidation), antibiotic stress response (**Figure 6A.1**), or redox balance and free radical scavenging (**Figure 6A.2**) (Anantharaman et al., 2014; Gyaneshwar et al., 2005; Mahmoudabadi et al., 2017; Nambi et al., 2015; Pal et al., 2018; Roux et al., 2016; Xia et al., 2017). To benefit the virus directly, sulfide could be utilized for amino acid synthesis or protein function, such as for co-factor binding (e.g. metal ions) (**Figure 6A.3.1**), persulfidation of cysteine residues for signaling (**Figure 6A.3.2**), structural sulfide bridge formation (**Figure 6A.3.3**), iron-sulfur cluster formation (**Figure 6A.3.4**) or for viral structural proteins in virion assembly (**Figure 6A.4**) (Peng et al., 2017; Tam et al., 2013). Furthermore, thiol modification of nucleic acids (i.e. dsDNA, tRNA and sRNA) could provide an avenue for responding to stresses (**Figure 6A.5**) or regulating gene expression for the virus or host (**Figure 6A.6**) (Damon et al., 2015; Hsu et al., 1967; Lira et al., 2018; Peng et al., 2017; Shimizu et al., 2017; Yang et al., 2017). Another method of nucleic acid modification that viruses may rely on is dsDNA recombination or integration (**Figure 6A.7.1**), or dsDNA repair (**Figure 6A.7.2**) which can be enabled by essential thiol components of enzymes (Jessop et al., 2000; Kessler, 2006; Yeeles et al., 2009). Sulfide may even be a key component in the ability of viruses to effectively lyse their host (**Figure 6A.8**) (Propst-Ricciuti, 1976).

**Figure 6.**
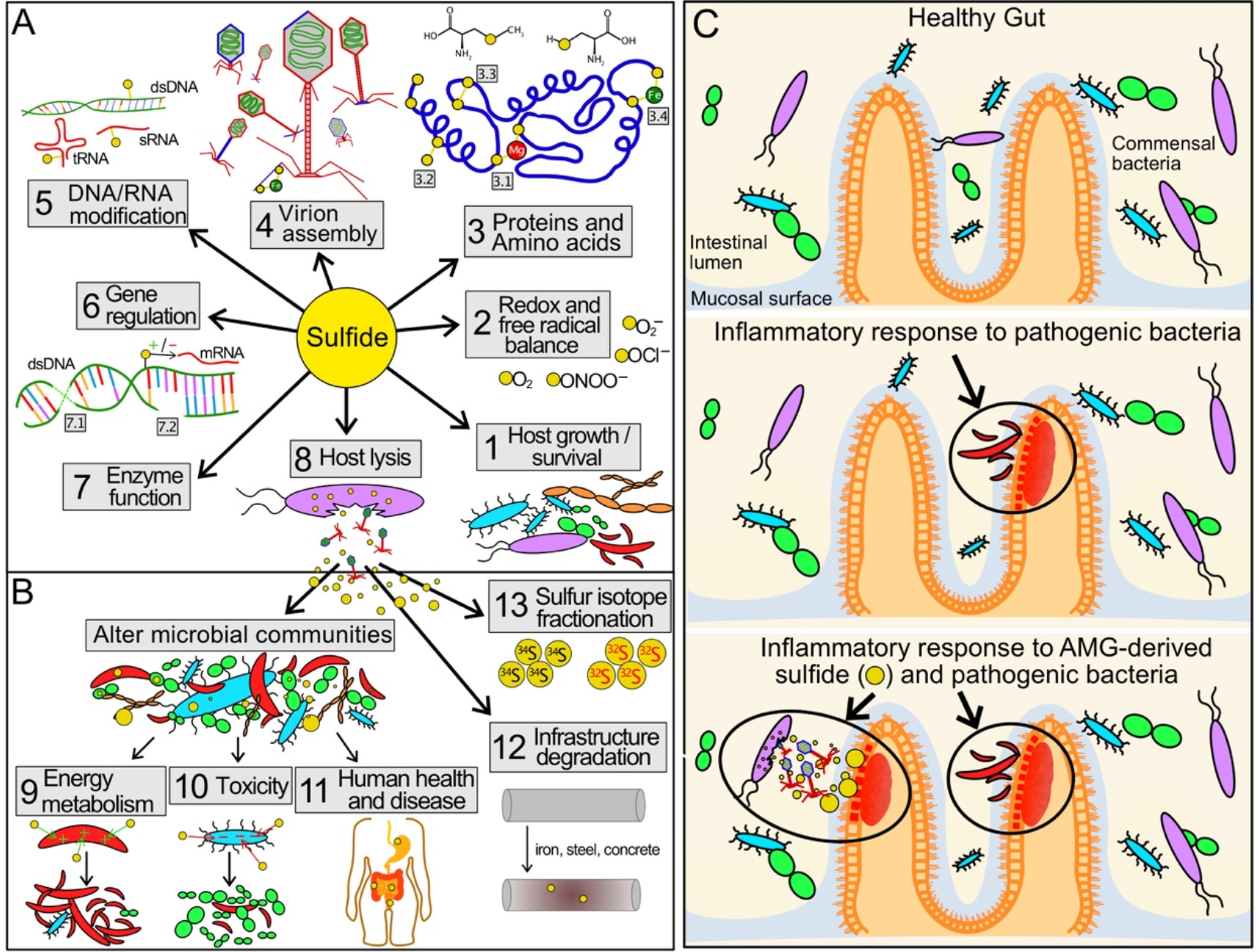
Virus-driven production of sulfide and its effects on human health, viral fitness and microbial communities. (A) Mechanisms by which sulfide could benefit viral fitness and (B) effect microbial communities, human health and environmental conditions. (C) Proposed impact of viral driven production of sulfide, in conjunction with activity of pathogenic bacteria, on inflammation in the gastrointestinal tract and its implications in IBD and CRC.

However, due to the diversity of functions encoded by AMGs (e.g. degradation of organosulfur compounds directly into sulfide or sulfite, manipulation of organosulfur compound forms or fixing sulfur) it is likely that host physiology and local environmental conditions drive their acquisition and function. Regardless of the utility of AMGs employed by individual viruses, the eventual lysis and release of virus-derived sulfide or virus-influenced sulfur chemistry could have significant impacts on the surrounding environment and local microbial communities (**Figure 6B**). Increased sulfide concentrations could either enhance the growth of sulfide oxidizing organisms (**Figure 6B.9**) or act as a toxin to inhibit the growth of others (**Figure 6B.10**) (Pal et al., 2018). Likewise, in both environmental and human systems, intracellular content released through viral lysis could alter nutrient availability and sulfide concentrations in the microbial community (**Figure 6B.11**) or lead to the degradation of iron, steel and concrete in infrastructure (**Figure 6B.12**).

In humans, balancing organic and inorganic sulfur concentrations is pivotal to both the health of the gastrointestinal tract and the resident microbiota (Yin et al., 2016), and our evidence suggests that viruses may interfere with this equilibrium. Moreover, dozens of microbial species have been linked to accumulation of sulfide within the human gut via the degradation of organosulfur compounds (e.g. cysteine and taurine) and implicated in CRC and IBD (Carbonero et al., 2012; Guo et al., 2016), but the role of viruses in facilitating or upregulating these processes is unknown. Specifically, virus-mediated sulfide production could accelerate the development of sulfide-associated gastrointestinal disorders such as colitis, IBD and CRC (**Figure 6C**).

Our discovery of AMGs for organosulfur metabolism and sulfide production also has widespread ramifications for interpreting Earth history (**Figure 6B.13**). Sulfur isotope fractionation (^34^S/^32^S) analysis is widely used to interpret geological records and estimate rates of microbial processes such as sulfate reduction (Habicht and Canfield, 1997; Sim et al., 2019; Thode et al., 1953). Microbial assimilatory sulfate reduction and viral auxiliary metabolism have been ignored as contributors to fractionation in the environment, mainly because sulfide is incorporated into organosulfur compounds instead of being exported into the environment as it is in dissimilatory reactions. As a result, assimilatory fractionation appears to be negligible (~3‰), whereas dissimilatory fractionation is frequently measured closer to 47‰ (Chambers and Trudinger, 1979; Kaplan and Rittenberg, 1964). Without the incorporation of sulfide into organosulfur compounds, assimilatory sulfite to sulfide reduction fractionates up to 36-42‰ in *Salmonella*, *Clostridium* and *Bacillus* species (Chambers and Trudinger, 1979). We propose that virus-mediated sulfide production can directly impact the observed fraction of ^32^S-enriched sulfide at scales relevant to dissimilatory sulfate reduction.

Overall, the global distribution and diversity of viruses encoding organosulfur transforming AMGs represents a novel and so-far unexplored cog in the global organic and inorganic sulfur cycles. By modulating organic and inorganic sulfur compound concentrations, viruses likely play important roles in infrastructure degradation, human disease and ecosystem health. Beyond viral organosulfur metabolism, this study serves as a model for elucidating the impacts of virus-driven degradation of amino acids, whose fate is an important driver in human health and biotechnology and associated with ecosystem services in agriculture.

## Acknowledgments

We also thank Anna-Louise Reysenbach and Katherine D. McMahon for helpful discussions and suggestions.

## Funding

We thank the University of Wisconsin - Office of the Vice Chancellor for Research and Graduate Education, University of Wisconsin – Department of Bacteriology, and University of Wisconsin - College of Agriculture and Life Sciences for their support. AJP was supported by the Ministry of Culture and Science of North Rhine-Westphalia (Nachwuchsgruppe “Dr. Alexander Probst”) and the NOVAC project of the German Science Foundation (grant number DFG PR1603/2-1). The work conducted by the U.S. Department of Energy Joint Genome Institute is supported by the Office of Science of the U.S. Department of Energy under contract no. DE-AC02-05CH11231.

## Author contributions

K.K. and K.A. designed the study. K.K., E.Z., P.H., and A.M.B. performed host-virus experiments. K.K. and K.A. conducted bioinformatic and metabolic analyses. K.K. and A.L. performed metatranscriptomic analyses. K.K. and K.A drafted the manuscript. All authors reviewed the results and approved the manuscript.

## Competing interests

The authors declare no competing interests.

## STAR Methods

### LEAD CONTACT AND MATERIALS AVAILABILITY

#### Lead Contact

Further information and any resources requests should be directed to and will be fulfilled by the lead contact Karthik Anantharaman.

#### Materials Availability

The recombinant phage line generated in this study is available upon request.

#### Data and Code Availability

All sequences used in this study are publicly available and can be found at their original sources. The genomic and protein sequences of viruses highlighted in this study and respective AMG protein sequences identified can be found at https://github.com/AnantharamanLab/Kieft_et_al_2020_organosulfur_AMGs.

### KEY RESOURCES TABLE

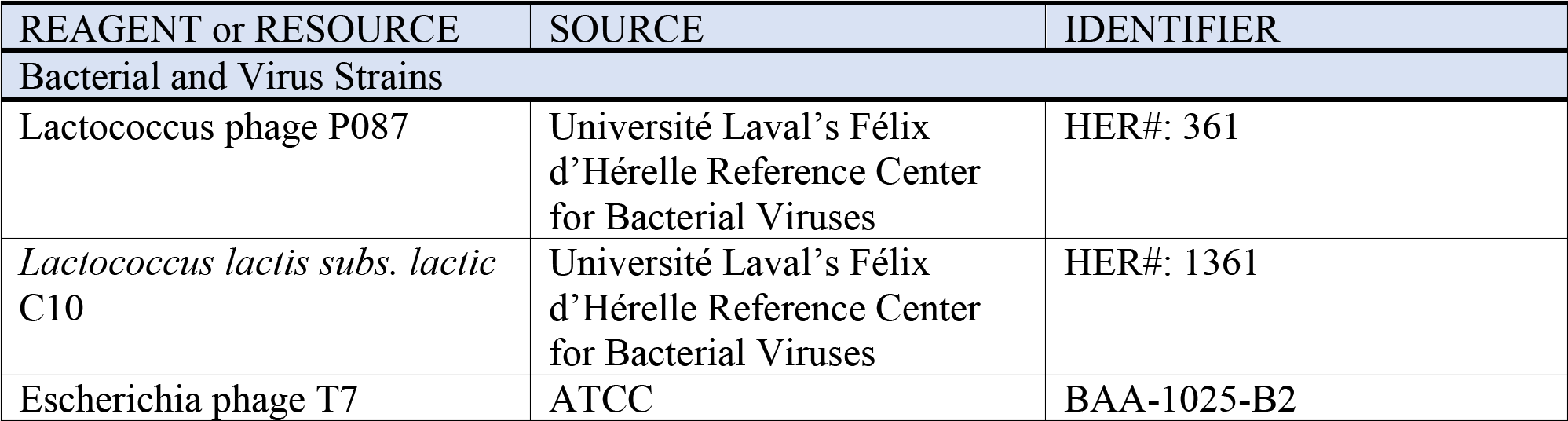
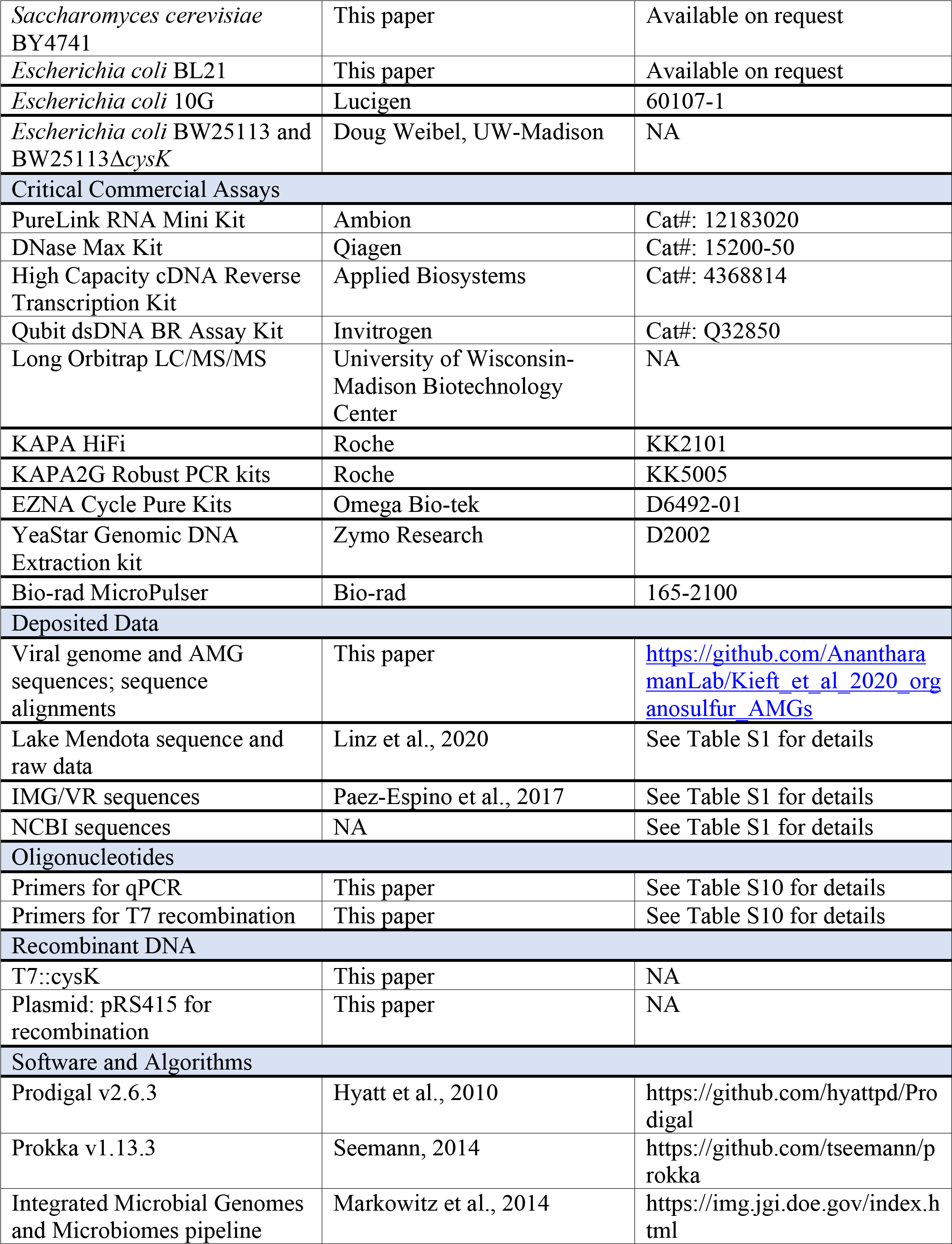
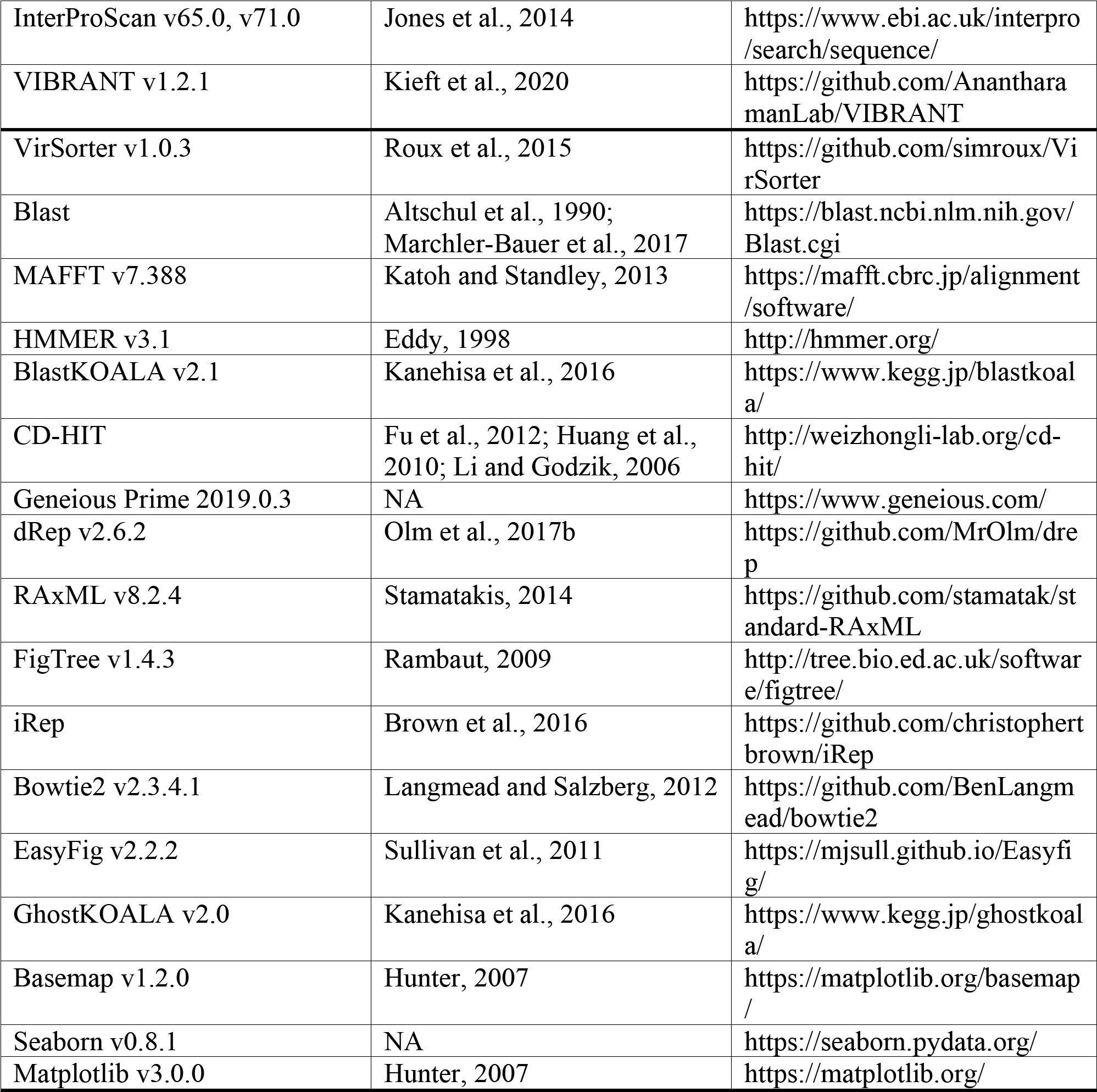

### EXPERIMENTAL MODEL AND SUBJECT DETAILS

#### *Lactococcus* system growth conditions

*Lactococcus lactis subs. lactis* C10 and Lactococcus phage P087 were obtained from Université Laval’s Félix d’Hérelle Reference Center for Bacterial Viruses (Canada, www.phage.ulaval.ca). *L. lactis* C10 was grown without agitation at 30°C in M17 broth (Oxoid) supplemented with 0.5% glucose (GM17). Infections were supplemented with 10mM CaCl_2_ and incubated without agitation at room temperature.

#### T7 system growth conditions

T7 phage was obtained from ATCC (ATCC^®^ BAA-1025-B2). *Saccharomyces cerevisiae* BY4741 and *E. coli* BL21 are lab stocks, *E. coli* 10G is a highly competent DH10B derivative (Durfee et al., 2008) originally obtained from Lucigen (60107-1). *E. coli* BW25113 and BW25113Δ*cysK* were obtained from Doug Weibel (University of Wisconsin, Madison).

All bacterial hosts were grown in and plated on LB media (1% Tryptone, 0.5% Yeast Extract, 1% NaCl in dH_2_O, plates additionally contain 1.5% agar, while top agar contained 0.5% agar) and LB media was used for all experimentation. All incubations of bacterial cultures were performed at 37°C, with liquid cultures shaking at 200-250 rpm unless otherwise specified. Bacterial hosts were streaked on appropriate LB plates and stored at 4°C. *S. cerevisiae* BY4741 was grown on YPD (2% peptone, 1% yeast extract, 2% glucose in dH_2_O, plates additionally contain 2.4% agar), after Yeast Artificial Chromosomes (YAC) transformation *S. cerevisiae* BY4741 was grown on SD-Leu (0.17% yeast nitrogen base, 0.5% ammonium sulfate, 0.162% amino acids – Leucine [Sigma Y1376], 2% glucose in dH_2_0, plates additionally contain 2% agar). All incubations of *S. cerevisiae* were performed at 30°C, with liquid cultures shaking at 200-250 rpm. *S. cerevisiae* BY4741 was streaked on YPD or SD-Leu plates as appropriate and stored at 4°C.

T7 phage was propagated using *E. coli* BL21 after initial receipt from ATCC and then as described on various hosts in methods. All phage experiments were performing using LB and culture conditions as described for bacterial hosts. Phages were stored in LB at 4°C. For long term storage all microbes were stored as liquid samples at −80°C in 10% glycerol, 90% relevant media. SOC (2% tryptone, 0.5% yeast extract, 0.2% 5M NaCl, 0.25% 1M KCL, 1% 1M MgCl_2_, 1% 1M MgSO_4_, 2% 1M glucose in dH_2_O) was used to recover host and phages after transformation.

For infection experiments, stationary phase cultures were created by growing bacteria overnight (totaling ~20-30 hours of incubation) at 37°C. Exponential phase culture consisted of stationary culture diluted 1:20 in LB then incubated at 37°C until an OD_600_ of ~0.4-0.8 was reached, typically after 40 minutes. Phage lysate was purified by centrifuging phage lysate at 16g, then filtering supernatant through a 0.22 μM filter. To establish titer, phage samples were serially diluted (1:10 or 1:100 dilutions made to 1 mL in 1.5 mL microcentrifuge tubes) in LB to a 10^-8^ dilution for titering by spot assay. Spot assays were performed by mixing 250 μl of relevant bacterial host in the stationary phase with 3.5 mL of 0.5% top agar, briefly vortexing, then plating on LB plates pre-warmed to 37°C. After plates solidified (typically ~5 minutes), 1.5 μl of each dilution of phage sample was spotted in series on the plate. Plates were incubated and checked every 2-4 hours or overnight (~20-30 hours) to establish a preliminary titer. MOI was estimated by calculated by dividing phage titer by estimated bacterial concentration.

### METHOD DETAILS

#### Identification of viral genomes

A total of 125,842 viral genomes from the Integrated Microbial Genomes/Virus (IMG/VR) (Paez-Espino et al., 2017) v1 database were used for analysis (accessed October 2017). Only publicly available genomes >5kb analyzed by Paez-Espino *et al*. (2016) were used in this study (Paez-Espino et al., 2016). Open reading frames were predicted using Prodigal with default parameters (v2.6.3) (Hyatt et al., 2010). All viral genomes were annotated using a combination of Prokka (v1.13.3) (Seemann, 2014), Integrated Microbial Genomes and Microbiomes pipeline (Markowitz et al., 2014), and InterProScan (v65.0) (Jones et al., 2014). Contigs with a high ratio of bacterial to viral protein annotations were manually identified and discarded. Contigs were further validated and annotated using a combination of VIBRANT (v1.2.1) and VirSorter (v1.0.3, virome database, categories 1, 2, 4, 5) (Kieft et al., 2020; Roux et al., 2015). All viral genomes encoding AMGs were manually inspected. Additional viral genomes were identified on the National Center for Biotechnology Information (NCBI) RefSeq (Brister et al., 2015; O’Leary et al., 2016; Tatusova et al., 2016) or Genbank database (Clark et al., 2016) (accessed Jan 2019) by querying viral genomes for AMGs of interest by blastp domain analysis (Altschul et al., 1990; Marchler-Bauer et al., 2017). Approximately 9,500 genomes corresponding to the viral classification *Caudovirales* were searched. VIBRANT and VirSorter were used to identify viruses >5kb from Lake Mendota, WI.

#### AMG identification and annotation

In-house hidden Markov model (HMM) profiles were built corresponding to the Kyoto Encyclopedia of Genes and Genomes (KEGG) pathway of organosulfur Metabolism as well as Cysteine and Methionine Metabolism (accessed December 2018) (Kanehisa and Goto, 2000). The two pathways’ KEGG Orthology (KO) numbers (189 total) were used to access corresponding proteins from the UniProt database (release 2018_11) (UniProt Consortium, 2018). The resulting proteins were aligned with MAFFT (v7.388, default parameters) (Katoh and Standley, 2013) and HMM profiles were built using hmmbuild (HMMER v3.1, default parameters) (Eddy, 1998). HMM profiles for CysC and CysH were built in the same manner, except manually verified viral CysC and CysH sequences, respectively, were added to the alignment for robustness. Hmmsearch (HMMER v3.1, evalue < 1e-5) was used to scan proteins on viral genomes. Proteins identified by the in-house HMM profiles were uploaded to the KEGG BlastKOALA server (v2.1) (Kanehisa et al., 2016) and queried under “prokaryotes” taxonomy and “genus prokaryotes” database for best hit annotations. Proteins annotated according to the original 189 KO numbers were selected for further verification. Manual verification of several representatives of each KO number (i.e. protein family) was done to curate the results using blastp (NCBI non-redundant database, accessed Jan 2019) and InterProScan (v71.0) to check for the presence of all expected conserved domains. Individual proteins and protein families of irrelevance or incorrect annotation were removed.

#### Sequence alignment and *dN/dS* analysis

Alignment of CysH, CysK, CysC, TauD and MetK sequences was performed using MAFFT (v7.388, default parameters). For *cysH-encoding* genomes identified from NCBI, all viral sequences were used. Host genomes were scanned, by annotation and blastp domain analysis, for multiple copies of *cysH* and all those identified were used, along with non-host bacterial sequences that were found to be highly similar to viral sequences according to pairwise identity. For the remaining alignments, all viral AMG protein sequences that shared at least 95% pairwise identity were restricted to one representative using CD-HIT (accessed Jan 2019) (Fu et al., 2012; Huang et al., 2010; Li and Godzik, 2006) and aligned. Viral CysK and CysH sequences were limited to lengths 200-330 and 117-600 amino acids, respectively. To obtain bacterial representatives, the majority consensus sequence of aligned viral proteins was queried against the NCBI RefSeq database by blastp (evalue < 1e-5). In order to ensure broad phylogenetic distribution of blastp results, the output was restricted to the top 500 hits from each of five phylogenetic groups based on NCBI categorization: [1] Proteobacteria, [2] Terrabacteria, [3] FCB superphlylum, [4] PVC superphylum and [5] a group containing all other phyla. The resulting sequences were manually limited to specific lengths to match viral sequences (CysC: 210-360, CysH: 150-600, CysK: 269-400, TauD: 314-400 amino acids, MetK: all) and reduced to one representative per 50% pairwise identity using CD-HIT. Viral and bacterial representatives were aligned together using MAFFT (default parameters) and gaps were stripped by 98%. The resulting alignments were used for phylogenetic analysis. Visualization of alignments was done using Geneious Prime 2019.0.3. For reference to full virus protein name and genome, see Table S1.

The AMGs for *cysK, cysC, cysD, cysH, tauD, msmA, metK, mtmN and luxS* were used to calculate *dN/dS* ratios. dRep (v2.6.2) was used to compare AMG sequences separately (dRep compare --SkipMash --S_algorithm goANI) and dnds_from_drep.py was used to calculate *dN/dS* ratios from the AMG pairs (Olm et al., 2017b). The *dN/dS* ratios were visualized with Seaborn (v0.8.1) and Matplotlib (v3.0.0).

#### Sequence phylogeny

Phylogenetic analysis was performed using protein alignments of CysH, CysK, CysC, TauD and MetK as described above. To infer phylogenetic relationships RAxML (v8.2.4) (Stamatakis, 2014) was used with the following parameters: raxmlHPC-PTHREADS -N 100 -f a -m PROTCATLG. Resulting best trees were used and rooted by manual identification of most distant (outgroup) taxa. Trees were visualized using FigTree (v1.4.3) (Rambaut, 2009).

#### Protein functional analysis

For domain and residue analysis, phylogenetic trees were used as a reference to select representative viral and bacterial sequences, which were then aligned using MAFFT (default parameters). Annotations of functional amino acid residues were labeled according to the Protein Data Bank (PDB, accessed January 2019) (Berman et al., 2000) with the following identification numbers: 4BZQ and 4BZP (CysC), 2GOY (CysH), 3ZEI (CysK), 3SWT (TauD), and 1RG9 (MetK). For alignments with no phylogenetic tree, up to five viral sequences and five PDB homologs (when available) were randomly selected for all AMGs with abundance of five or greater. The PDB sequences used for annotation were added to the alignment. N- and C-terminal ends of protein alignments were manually removed for clarity and gaps were stripped by 90% (for alignments with phylogenetic trees) or 80% (for all others). Residues were highlighted according to 85% pairwise identity between sequences, excluding sequence gaps. An identity graph, generated by Geneious, was fitted to the alignment to visualize pairwise identity of 100% (green), 99-30% (yellow) and 29-0% (red).

#### Protein Reactions

Enzymatic reactions, diagrams and pathways were created by referencing KEGG and MetaCyc (v22.6) (Caspi et al., 2012) annotations.

#### Viral transcriptomics and growth rates

Publicly available metatranscriptomic data from Lake Mendota, WI was assessed for AMGs by querying annotation names (Linz et al., 2020). This gene expression data comprises a two-day time series and is accompanied by metagenomic assemblies (IMG Taxon Object IDs 3300013004 and 3300013005). Metatranscriptomic reads were mapped to a custom, non-redundant database of freshwater reference data, including the metagenome assemblies; annotations in this study are derived from the annotations of the reference database. We used read counts normalized to transcripts per liter as the input for our study, and we searched for AMGs in the metagenomic assemblies as described above.

The growth rate of the *cysC*-encoding Lake Mendota virus was identified using index of replication (iRep) with default parameters (Brown et al., 2016). Metagenomic assembly reads used for iRep are available on IMG under the Taxon Object ID 3300013005. Reads were mapped to the viral genome using Bowtie2 (v2.3.4.1) (Langmead and Salzberg, 2012). GC-skew to indicate rolling circle replication of the viral genome was likewise completed using the iRep toolkit.

#### Virus growth and fitness assay

Approximately 10^8^ plaque forming units (PFUs) of Lactococcus phage P087 (approximate multiplicity of infection (MOI) of 1) were used to infect 1mL of *L. lactis* C10 which had been brought to an optical density (OD_600_) of approximately 0.15 in GM17 broth. For fitness experiments, either vehicle control (water), 10μM Na2S or 100μM Na2S was supplemented to the media at time of infection. Infections were incubated without agitation at room temperature for approximately three hours. Additional cultures of uninfected *L. lactis* C10 with all other variables identical were measured for growth at the endpoint of infections using OD_600_. To end infections, *L. lactis* C10 were spun out of solution at 10,000 rcf and the supernatant (i.e viral fraction) was removed and cooled to 4°C. Plaque assays were done using the standard double agar method (Lillehaug, 1997) with diluted viral fraction and *L. lactis* C10 brought to high concentration. A 1% bottom agar and 0.4% top agar of GM17 were used, both supplemented with 0.5% glycine and 10mM CaCl_2_.

#### Virus and host *cysK* qPCR assay

An overnight culture of *L. lactis* C10 was diluted in GM17 broth to OD 0.08 and grown at 30°C for ~2 hours until OD reached 0.15. In a batch culture 10mM CaCl_2_ was added. Two different conditions were assayed, each in duplicate: (1) *L. lactis* C10 control and (2) *L. lactis* C10 plus Lactococcus phage P087. For infection conditions, Lactococcus phage P087 was added at a MOI of 1 (time 0 minutes). RNA was extracted using the PureLink RNA Mini Kit (Ambion) from 500μL of the cellular fraction at 15, 60 and 120 minutes post-infection. RNA was then treated with DNase with the DNase Max Kit (Qiagen) and converted to cDNA using the High Capacity cDNA Reverse Transcription Kit (Applied Biosystems). qPCR of viral and host *cysK* was performed using *Power* SYBR Green PCR Master Mix (Applied Biosystems) with 7ng of cDNA template and the following primer sets (IDT): *L. lactis* C10 forward (CCTTCGTTGGCTCTGCTTTG), *L. lactis* C10 reverse (TGGCATCATCTCCTTTGACCC), Lactococcus phage P087 forward (CAGAAACTATCGGAAACACACCAC), and Lactococcus phage P087 reverse (TTGAGTGAATGACCTGCTCCA) (**Table S10**). The concentration of template cDNA was measured with the Qubit dsDNA BR Assay Kit (Invitrogen). The viral and host *cysK* sequences were sufficiently dissimilar in sequence identity (<60% at the protein level) to allow for accurate distinction by qPCR and the primers selected.

#### Mass spectrometry and protein identification

*L. lactis* C10 was grown without agitation at 30°C in modified M17 broth supplemented with 0.5% glucose (mGM17). mGM17 was made by adding 1.25g glucose, 0.625g tryptone, 1.25g peptone, 0.125g yeast extract, 0.125g ascorbic acid, 0.0626g anhydrous magnesium sulfate and 4.75g disodium glycerophosphate to 250mL deionized water. Approximately 10^8^ PFUs of Lactococcus phage P087 were used to infect 3mL of *L. lactis* C10 which had been brought to OD_600_ of approximately 0.15 and supplemented with 10mM CaCl_2_. Infections proceeded to complete lysis without agitation at room temperature for approximately three hours. To end the infection, *L. lactis* C10 were spun out of solution at 10,000 rcf and the supernatant was removed and stored at 4°C. The supernatant was size fractionated by filtration for the 100kDa to 10kDa size fraction before trypsin solution digestion and analysis by Long Orbitrap LC/MS/MS (University of Wisconsin-Madison Biotechnology Center).

#### Genome organization and comparisons

Genome organization was visualized using Geneious Prime. Genes were manually colored by referencing functions according to NCBI RefSeq or Genbank annotation, or blastp search._Viral genomes in genbank format were compared and visualized with EasyFig (v2.2.2) (Sullivan et al., 2011) using the tblastx function. Only tblastx (v2.8.1+) hits with percent identities greater than 30% and e-values less than 0.001 are shown. Remaining analysis parameters were set to default. Circular sequences were visualized linearly for ease of comparison.

#### Geographical distributions

IMG Taxon Object ID numbers were used to identify global coordinates of studies in which AMGs were identified. Coordinates were mapped using Matplotlib’s Basemap (v1.2.0) (Hunter, 2007). Human studies were excluded from coordinate maps.

#### Host classification

GhostKOALA (v2.0) (Kanehisa et al., 2016) with the “genus prokaryotes” database was used to query all 3,794 AMG-encoded proteins identified from IMG/VR derived viruses (3,421 annotated and used for taxonomy). To benchmark accuracy of the analysis, all 282 AMG-encoded proteins identified from NCBI-derived viruses with known hosts were queried in the same manner (278 were annotated and used for taxonomy) and compared to the taxonomy of hosts.

#### T7 recombination: cloning

All primers can be found in **Table S10**. PCR was performed using KAPA HiFi (Roche) for all experiments with the exception of multiplex PCR for screening Yeast Artificial Chromosomes (YACs), which was performed using KAPA2G Robust PCR kits (Roche). DNA purification was performed using EZNA Cycle Pure Kits (Omega Bio-tek) using the centrifugation protocol. YAC extraction was performed using YeaStar Genomic DNA Extraction kits (Zymo Research). All cloning was performed according to manufacturer documentation except where noted in methods. PCR reactions using phage as template use 1 μl of undiluted phage stock, with extension of the 95°C denaturation step to 5 minutes.

Electroporation of YACs was performed using a Bio-rad MicroPulser (165-2100), Ec2 setting (2 mm cuvette, 2.5 kV, 1 pulse) using 50 μl competent cells and 2 μl YAC DNA for transformation. Electroporated cells were immediately recovered with 950 μl SOC, then incubated at 37°C for 1 to 1.5 hours and plated or grown in Lb.

*E. coli* 10G competent cells were made by adding 8 mL overnight 10G cells to 192 mL SOC (with antibiotics as necessary) and incubating at 21°C and 200 rpm until ~OD_600_ of 0.4 as determined using an Agilent Cary 60 UV-Vis Spectrometer using manufacturer documentation (actual incubation time varies based on antibiotic, typically overnight). Cells are centrifuged at 4°C, 800-1000g for 20 minutes, the supernatant is discarded, and cells are resuspended in 50 mL 10% glycerol. Centrifugation and washing are repeated three times, then cells are resuspended in a final volume of ~1 mL 10% glycerol and are aliquoted and stored at −80°C. Cells are competent for plasmid and YACs. All primers used in experiments in this publication are listed in supplemental.

#### T7 recombination: engineering T7 with *cysK*

Phages were assembled using YAC rebooting (Ando et al., 2015; Jaschke et al., 2012), which requires yeast transformation of relevant DNA segments, created as follows. A prs415 yeast centromere plasmid was split into three segments by PCR, separating the centromere and leucine selection marker, which partially limits recircularization and improved assembly efficiency (Kuijpers et al., 2013). Wildtype T7 segments were made by PCR using wildtype T7 as template. *CysK* segments were made by colony PCR of BW25113. *CysK* was inserted into two locations to create two phage constructs. The first location was replacement of gp1.7 to establish *CysK* in early Class II genes. This insertion causes a two amino acid extension (YE) of the immediate 5’ gene gp1.6 that was not anticipated to have an effect on phage viability. The second location was inserted adjacent to gp6.3 to establish *CysK* in early class III genes and leverages a copy of phage promoter phi6.5 for expression.

DNA parts were combined together (0.1 pmol/segment) and transformed into *S. cerevisiae* BY4741 using a high efficiency yeast transformation protocol (Daniel Gietz and Woods, 2002) using SD-Leu selection. After 2-3 days colonies were picked and directly assayed by multiplex colony PCR to assay assembly. Multiplex PCR interrogated junctions in the YAC construct and was an effective way of distinguishing correctly assembled YACs. Correctly assembled YACs were purified and transformed into *E. coli* 10G cells and these cultures incubated until lysis, after which phages were purified to create the initial phage stock.

#### T7 recombination: passaging and AMG retention

Either T7Δ1.7::cysK or T7::cysK phages were added to 5 mL exponential phase BW25113 or BW25113Δ*cysK* at an estimated MOI of 10^-4^ to allow for an estimated three phage passages. After the culture had fully lysed, typically ~1 hour and 30 minutes, lysate was purified and then the titer established by spot assay. This process was then repeated twice for a total of an estimated 9 phage passages assuming at least 100 phage progeny per host. Phage lysate from the final passage was used as template for sequencing to determine if the *cysK* insert remained as the consensus sequence in the phage population. The entire process was repeated in biological triplicate for both host and phage combinations.

### QUANTIFICATION AND STATISTICAL ANALYSIS

#### Virus growth and fitness

The number of resulting plaques from the growth and fitness assays were normalized to 100% of controls for each experiment. Three independent experiments with three infection replicates and two growth replicates each was performed. Further information of experiments can be found in Method Details below.

#### Virus and host *cysK* qPCR

For each replicate of the two conditions assayed both primer sets were used for qPCR. To analyze the qPCR results, the Cq readings were averaged between the three replicates for each treatment at each timepoint to obtain a single datapoint per treatment:primer pair per timepoint, termed *average Cq*. Using time point zero for the uninfected *L. lactis* C10 condition with *L. lactis* C10 *cysK* primers as the baseline *control*, delta-delta-Cq values were calculated by subtracting the *control* value from the *average Cq* values. This result calculates the expression of *L. lactis* C10 *cysK* at time point zero to be normalized to zero (delta-delta-Cq of zero). Finally, all delta-delta-Cq values were transformed using the formula 2^-(delta-delta-Cq)^ (Livak and Schmittgen, 2001). All raw Cq values and normalized results, including equations, can be found in **Table S6**. Further information of experiments can be found in Method Details below.

## Supplemental Information

**Table S1** (Separate File). **Metadata for all viruses identified in this study, related to Figures 1–5.** List of all viral genomes with associated metadata: database origin, AMG name, KEGG orthology number, name of protein product, viral AMG and genome name, genome length, shorthand name if appended in study, environment and geographical information, cultivation status, accession numbers, virus classification, host classification and human gut association, and isolation location.

**Table S2** (Separate File). **Abundance of each AMG per environment or human microbiome, related to Figure 2A.** Only AMGs with IMG/VR environmental categories are shown.

**Table S3** (Separate File). **Number of AMGs and IMG studies per environment classification, related to Figure 2B.** Only AMGs with IMG/VR environmental categories are shown.

**Table S4** (Separate File). **Richness of viruses encoding AMGs per environment, related to Figure 2A, B.** Shown is the number of IMG/VR non-redundant viral genomes encoding at least one AMG per environment, the total number of viruses in the IMG/VR dataset per environment and the percentage of viruses per environment that encode at least one AMG.

**Table S5** (Separate File). **Expression of viral AMGs in Lake Mendota, WI, related to Figure 6.** Viral genomes expressing the indicated AMG over two days (timepoint). Expression is given as a measure of read counts normalized to an internal standard.

**Table S6** (Separate File). **qPCR raw and normalized results of *cysK* transcript relative abundance for *L. lactis* C10 and phage P087, related to Figure 3.** The raw Cq results for each condition and primer set are provided, in addition to the normalization equations and 2^-ddCq^ results used to generate Figure S5.

**Table S7** (Separate File). **Complete profile of proteins identified by untargeted proteomics between 100kDa and 10kDa, related to Figure 3**. The best-hit identification of peptides according to NCBI databases, their associated accession number, calculated molecular weight and spectral count. Lactococcus phage P087 and *L. lactis* CysK were identified to be 34^th^ and 57^th^ most abundant, respectively.

**Table S8** (Separate File). **Full results of AMG protein identity search, related to Figures 2 and 6.** Tabular formatted blastp output of AMG queries against the NCBI non-redundant (nr) database.

**Table S9** (Separate File). **List of viral genomes containing multiple AMGs, related to Figures 1, 2, 4 and 5**. Shown are the viral gene and genome name corresponding to each encoded AMG.

**Table S10** (Separate File). **List of all sequence primers used in this study, related to Figure 3**. Primers are listed for the virus and host *cysK* qPCR assay as well as for T7 recombination.

**Additional Data File S1** (Separate File)**. Amino acid sequences of each of the 4,103 AMGs identified in this study.** Each file is named with the three- or four-letter protein ID of the AMG followed by the respective KEGG orthology number.

**Additional Data File S2** (Separate File)**. Amino acid sequence alignments, in FASTA format, respective to Figures S2 and S4.**

**Additional Data File S3** (Separate File)**. Full genome sequence and Bowtie2 alignment of the *cysC*-encoding Lake Mendota virus.** The genome sequence and alignment files were used for the generation of Figure S3.

**Figure S1. Distribution of individual AMGs in the environment, related to Figure 2.** Global distribution of viral populations encoding (A) *cysC*, (B) *cysH*, (C) *cysK*, (D) *dcm*, (E) *metK*, (F) *tauD* or (G) *msmA* identified on the IMG/VR database, color coordinated by environment classification.

**Figure S2. Amino acid alignments of proteins encoded by AMGs, related to Figure 1 and Table 1.** Alignment of representative viral and bacterial sequences for all AMGs with abundances greater than five. Viruses are indicated by the preface “Phage” followed by their respective IMG Taxon Object ID number. See Table S1 for full genome names. Refer to Figurer S4 for phylogeny of the represented sequences for A-E. Highlighted amino acids indicate conservation in >85% of sequences shown. Black boxes indicate amino acid residues that are biochemically verified as functional according to the given PDB reference sequence.

**Figure S3. *dN/dS* ratio calculations for *cysK, cysC, cysD, cysH, tauD, msmA, metK, mtmN and luxS* AMG pairs, related to Figures 1, 4 and 5.** Each data point overlaid on the standard box plot represents a single AMG pair. The figure was generated using seaborn and matplotlib.

**Figure S4. Genome and growth statistics of a complete virus identified to express *cysC* in Lake Mendota, WI, related to Figure 6.** The (A) GC-skew and (B) index of replication for a complete, circular virus identified in Lake Mendota, WI. GC-skew and replication statistics indicate that the virus was actively replicating at time of sampling and likely undergoes rolling circle replication.

**Figure S5. qPCR results of *cysK* transcript relative abundance for *L. lactis* C10 and phage P087, related to Figure 3.** Relative transcript abundance is provided in the normalized 2^-ddCq^ metric. Control is *L. lactis* C10 (host) alone and infection includes host plus phage P087. The times shown are ti (15 minutes), t2 (60 minutes) and t3 (120 minutes).

**Figure S6. Phylogeny of viral AMG encoded protein sequences, related to Figures 1, 4 and 5**. (A) Phylogenetic tree for CysH of complete viruses with known bacterial hosts. Viruses are in red and bacteria are in black, and proteins with an additional DUF4440 domain are highlighted in blue. Bacteria with multiple copies of *cysH* are appended with a letter (“A” through “D”). Refer to Table S1 for virus-host associations. Also shown are phylogenetic trees of uncultivated viruses with bacterial homologs and select cultivated viruses for (B) CysH (red and green highlighting refers to putative virus-host associations; compressed blue clade contains 36 viruses and 89 bacteria from several phyla), (C) CysC, (D) CysK (yellow highlighting refers to known virus-host associations), (E) TauD and (F) MetK. For trees (B-F) colored names refer to viruses (black), Bacteroidetes and other members of the FCB superphylum (red), Cyanobacteria (cyan), Verrucomicrobia and Planctomycetes and other members of the PVC superphylum (purple), Actinobacteria (yellow), and all other phyla in orange. For all trees, bootstrap values greater than 60 are shown, orange highlighting indicates respective genomes used for comparative genomics (see Figures 4 and S6), and arrows indicate sequences used for protein alignments (see Figure S2).

**Figure S7. Taxonomic classification of AMGs, related to Figure 2**. Inferred taxonomic classification at the phylum level of AMG-encoded protein sequences on NCBI-derived (A) viruses and (B) taxonomy of their known hosts, showing similar proportionality. (C) Inferred taxonomic classification at the phylum level of AMG-encoded protein sequences on IMG/VR-derived viruses.

**Figure S8. Genome comparisons of viruses encoding AMGs, related to Figure 4**. Comparisons of (A) incomplete viruses and complete *Streptococcus pneumoniae* viruses encoding *dcm* (pink), (B) incomplete viruses encoding *metK* (pink), (C) complete *Bacillus cereus* viruses encoding *cysH* (pink), and (D) complete *Streptococcus suis* viruses encoding *metK* (yellow) and *dcm* (pink). For all comparisons, predicted open readings frames are annotated by dark blue arrows and genomes are connected with lines according to tblastx identity. Refer to Figure S6 for phylogeny of AMGs for (B) and (C) (orange highlighting).

**Figure S9. Genome organization of *cysH*-encoding viruses, related to Figure 5.** Organization of linear and circular complete viral genomes that encode *cysH*. Arrows indicate open reading frames and are annotated by general function: virion structural assembly (green), auxiliary metabolism and general functions (red), nucleotide metabolism and genome replication (blue), lysis (orange) and unknown function (yellow). Pink stars indicate the location of *cysH*. Refer to Table S1 for virus details.

## Notes

https://github.com/AnantharamanLab/Kieft_et_al_2020_organosulfur_AMGs

### Competing Interest Statement

The authors have declared no competing interest.

## References

Ahlgren, N.A., Fuchsman, C.A., Rocap, G., and Fuhrman, J.A. (2019). Discovery of several novel, widespread, and ecologically distinct marine Thaumarchaeota viruses that encode amoC nitrification genes. The ISME Journal 13, 618–631.

Almeida, A., Mitchell, A.L., Boland, M., Forster, S.C., Gloor, G.B., Tarkowska, A., Lawley, T.D., and Finn, R.D. (2019). A new genomic blueprint of the human gut microbiota. Nature 1.

Altschul, S.F., Gish, W., Miller, W., Myers, E.W., and Lipman, D.J. (1990). Basic local alignment search tool. J. Mol. Biol. 215, 403–410.

Anantharaman, K., Duhaime, M.B., Breier, J.A., Wendt, K.A., Toner, B.M., and Dick, G.J. (2014). Sulfur Oxidation Genes in Diverse Deep-Sea Viruses. Science 344, 757–760.

Anantharaman, K., Hausmann, B., Jungbluth, S.P., Kantor, R.S., Lavy, A., Warren, L.A., Rappé, M.S., Pester, M., Loy, A., Thomas, B.C., et al. (2018). Expanded diversity of microbial groups that shape the dissimilatory sulfur cycle. The ISME Journal 12, 1715.

Ando, H., Lemire, S., Pires, D.P., and Lu, T.K. (2015). Engineering Modular Viral Scaffolds for Targeted Bacterial Population Editing. Cels 1, 187–196.

Andreae, M.O. (1990). Ocean-atmosphere interactions in the global biogeochemical sulfur cycle. Marine Chemistry 30, 1–29.

Bakir, M.A., Kitahara, M., Sakamoto, M., Matsumoto, M., and Benno, Y. (2006). Bacteroides intestinalis sp. nov., isolated from human faeces. International Journal of Systematic and Evolutionary Microbiology 56, 151–154.

Berman, H.M., Westbrook, J., Feng, Z., Gilliland, G., Bhat, T.N., Weissig, H., Shindyalov, I.N., and Bourne, P.E. (2000). The Protein Data Bank. Nucleic Acids Res 28, 235–242.

Bragg, J.G., and Chisholm, S.W. (2008). Modeling the Fitness Consequences of a Cyanophage-Encoded Photosynthesis Gene. PLOS ONE 3, e3550.

Breitbart, M., Thompson, L., Suttle, C., and Sullivan, M. (2007). Exploring the Vast Diversity of Marine Viruses. Oceanography 20, 135–139.

Brister, J.R., Ako-Adjei, D., Bao, Y., and Blinkova, O. (2015). NCBI viral genomes resource. Nucleic Acids Res. 43, D571–577.

Brown, C.T., Olm, M.R., Thomas, B.C., and Banfield, J.F. (2016). Measurement of bacterial replication rates in microbial communities. Nature Biotechnology 34, 1256–1263.

Carbonero, F., Benefiel, A.C., Alizadeh-Ghamsari, A.H., and Gaskins, H.R. (2012). Microbial pathways in colonic sulfur metabolism and links with health and disease. Front Physiol 3.

Caspi, R., Altman, T., Dreher, K., Fulcher, C.A., Subhraveti, P., Keseler, I.M., Kothari, A., Krummenacker, M., Latendresse, M., Mueller, L.A., et al. (2012). The MetaCyc database of metabolic pathways and enzymes and the BioCyc collection of pathway/genome databases. Nucleic Acids Res. 40, D742–753.

Chambers, L.A., and Trudinger, P.A. (1979). Microbiological fractionation of stable sulfur isotopes: A review and critique. Geomicrobiology Journal 1, 249–293.

Chartron, J., Carroll, K.S., Shiau, C., Gao, H., Leary, J.A., Bertozzi, C.R., and Stout, C.D. (2006). Substrate Recognition, Protein Dynamics, and Iron-Sulfur Cluster in Pseudomonas aeruginosa Adenosine 5’-Phosphosulfate Reductase. J Mol Biol 364, 152–169.

Chen, L.-X., Méheust, R., Crits-Christoph, A., McMahon, K.D., Nelson, T.C., Warren, L.A., and Banfield, J.F. (2020). Large Freshwater Phages with the Potential to Augment Aerobic Methane Oxidation (Microbiology).

Chiku, T., Padovani, D., Zhu, W., Singh, S., Vitvitsky, V., and Banerjee, R. (2009). H2S Biogenesis by Human Cystathionine γ-Lyase Leads to the Novel Sulfur Metabolites Lanthionine and Homolanthionine and Is Responsive to the Grade of Hyperhomocysteinemia. J Biol Chem 284, 11601–11612.

Clark, K., Karsch-Mizrachi, I., Lipman, D.J., Ostell, J., and Sayers, E.W. (2016). GenBank. Nucleic Acids Res 44, D67–D72.

Costea, P.I., Coelho, L.P., Sunagawa, S., Munch, R., Huerta-Cepas, J., Forslund, K., Hildebrand, F., Kushugulova, A., Zeller, G., and Bork, P. (2017). Subspecies in the global human gut microbiome. Mol Syst Biol 13.

Curtis, M.M., Hu, Z., Klimko, C., Narayanan, S., Deberardinis, R., and Sperandio, V. (2014). The Gut Commensal Bacteroides thetaiotaomicron Exacerbates Enteric Infection through Modification of the Metabolic Landscape. Cell Host & Microbe 16, 759–769.

Damon, J.R., Pincus, D., and Ploegh, H.L. (2015). tRNA thiolation links translation to stress responses in Saccharomyces cerevisiae. Mol Biol Cell 26, 270–282.

Daniel Gietz, R., and Woods, R.A. (2002). Transformation of yeast by lithium acetate/single-stranded carrier DNA/polyethylene glycol method. In Methods in Enzymology, C. Guthrie, and G.R. Fink, eds. (Academic Press), pp. 87–96.

Devoto, A.E., Santini, J.M., Olm, M.R., Anantharaman, K., Munk, P., Tung, J., Archie, E.A., Turnbaugh, P.J., Seed, K.D., Blekhman, R., et al. (2019). Megaphages infect Prevotella and variants are widespread in gut microbiomes. Nature Microbiology 4, 693–700.

Durfee, T., Nelson, R., Baldwin, S., Plunkett, G., Burland, V., Mau, B., Petrosino, J.F., Qin, X., Muzny, D.M., Ayele, M., et al. (2008). The Complete Genome Sequence of Escherichia coli DH10B: Insights into the Biology of a Laboratory Workhorse. Journal of Bacteriology 190, 2597–2606.

Dutilh, B.E., Cassman, N., McNair, K., Sanchez, S.E., Silva, G.G.Z., Boling, L., Barr, J.J., Speth, D.R., Seguritan, V., Aziz, R.K., et al. (2014). A highly abundant bacteriophage discovered in the unknown sequences of human faecal metagenomes. Nature Communications 5, 4498.

Eddy, S.R. (1998). Profile hidden Markov models. Bioinformatics 14, 755–763.

Eeckhaut, V., Machiels, K., Perrier, C., Romero, C., Maes, S., Flahou, B., Steppe, M., Haesebrouck, F., Sas, B., Ducatelle, R., et al. (2013). Butyricicoccus pullicaecorum in inflammatory bowel disease. Gut 62, 1745–1752.

Fenner, L., Roux, V., Ananian, P., and Raoult, D. (2007). Alistipes finegoldii in Blood Cultures from Colon Cancer Patients. Emerg Infect Dis 13, 1260–1262.

Fike, D.A., Bradley, A.S., and Rose, C.V. (2015). Rethinking the Ancient Sulfur Cycle. Annual Review of Earth and Planetary Sciences 43, 593–622.

Fitzgerald, J.W. (1976). Sulfate ester formation and hydrolysis: a potentially important yet often ignored aspect of the sulfur cycle of aerobic soils. Bacteriol Rev 40, 698–721.

Fu, L., Niu, B., Zhu, Z., Wu, S., and Li, W. (2012). CD-HIT: accelerated for clustering the next-generation sequencing data. Bioinformatics 28, 3150–3152.

Gobler, C.J., Hutchins, D.A., Fisher, N.S., Cosper, E.M., and Saňudo-Wilhelmy, S.A. (1997). Release and bioavailability of C, N, P Se, and Fe following viral lysis of a marine chrysophyte. Limnology and Oceanography 42, 1492–1504.

Guo, F.-F., Yu, T.-C., Hong, J., and Fang, J.-Y. (2016). Emerging Roles of Hydrogen Sulfide in Inflammatory and Neoplastic Colonic Diseases. Front Physiol 7.

Gyaneshwar, P., Paliy, O., McAuliffe, J., Popham, D.L., Jordan, M.I., and Kustu, S. (2005). Sulfur and Nitrogen Limitation in Escherichia coli K-12: Specific Homeostatic Responses. Journal of Bacteriology 187, 1074–1090.

Habicht, K.S., and Canfield, D.E. (1997). Sulfur isotope fractionation during bacterial sulfate reduction in organic-rich sediments. Geochimica et Cosmochimica Acta 61, 5351–5361.

Henriques, A.C., and Marco, P.D. (2015). Methanesulfonate (MSA) Catabolic Genes from Marine and Estuarine Bacteria. PLOS ONE 10, e0125735.

Howard-Varona, C., Lindback, M.M., Bastien, G.E., Solonenko, N., Zayed, A.A., Jang, H., Andreopoulos, B., Brewer, H.M., Rio, T.G. del, Adkins, J.N., et al. (2020). Phage-specific metabolic reprogramming of virocells. ISME J 1–15.

Hsu, W.T., Foft, J.W., and Weiss, S.B. (1967). Effect of bacteriophage infection on the sulfur-labeling of sRNA. Proc Natl Acad Sci U S A 58, 2028–2035.

Huang, Y., Niu, B., Gao, Y., Fu, L., and Li, W. (2010). CD-HIT Suite: a web server for clustering and comparing biological sequences. Bioinformatics 26, 680–682.

Hugon, P., Ramasamy, D., Lagier, J.-C., Rivet, R., Couderc, C., Raoult, D., and Fournier, P.-E. (2013). Non contiguous-finished genome sequence and description of Alistipes obesi sp. nov. Stand Genomic Sci 7, 427–439.

Hunter, J.D. (2007). Matplotlib: A 2D graphics environment. Computing In Science & Engineering 9, 90–95.

Hurwitz, B.L., and U’Ren, J.M. (2016). Viral metabolic reprogramming in marine ecosystems. Current Opinion in Microbiology 31, 161–168.

Hurwitz, B.L., Hallam, S.J., and Sullivan, M.B. (2013). Metabolic reprogramming by viruses in the sunlit and dark ocean. Genome Biology 14, R123.

Hurwitz, B.L., Brum, J.R., and Sullivan, M.B. (2015). Depth-stratified functional and taxonomic niche specialization in the ‘core’ and ‘flexible’ Pacific Ocean Virome. ISME J 9, 472–484.

Hyatt, D., Chen, G.-L., LoCascio, P.F., Land, M.L., Larimer, F.W., and Hauser, L.J. (2010). Prodigal: prokaryotic gene recognition and translation initiation site identification. BMC Bioinformatics 11, 119.

Ishikawa, K., Mino, K., and Nakamura, T. (2010). New function and application of the cysteine synthase from archaea. Biochemical Engineering Journal 48, 315–322.

Jaschke, P.R., Lieberman, E.K., Rodriguez, J., Sierra, A., and Endy, D. (2012). A fully decompressed synthetic bacteriophage øX174 genome assembled and archived in yeast. Virology 434, 278–284.

Jessop, L., Bankhead, T., Wong, D., and Segall, A.M. (2000). The Amino Terminus of Bacteriophage □ Integrase Is Involved in Protein-Protein Interactions during Recombination. J. BACTERIOL. 182, 11.

Jiang, W., Wu, N., Wang, X., Chi, Y., Zhang, Y., Qiu, X., Hu, Y., Li, J., and Liu, Y. (2015). Dysbiosis gut microbiota associated with inflammation and impaired mucosal immune function in intestine of humans with non-alcoholic fatty liver disease. Scientific Reports 5, 8096.

Jiao, N., Herndl, G.J., Hansell, D.A., Benner, R., Kattner, G., Wilhelm, S.W., Kirchman, D.L., Weinbauer, M.G., Luo, T., Chen, F., et al. (2010). Microbial production of recalcitrant dissolved organic matter: long-term carbon storage in the global ocean. Nature Reviews Microbiology 8, 593–599.

Jones, P., Binns, D., Chang, H.-Y., Fraser, M., Li, W., McAnulla, C., McWilliam, H., Maslen, J., Mitchell, A., Nuka, G., et al. (2014). InterProScan 5: genome-scale protein function classification. Bioinformatics 30, 1236–1240.

Jover, L.F., Effler, T.C., Buchan, A., Wilhelm, S.W., and Weitz, J.S. (2014). The elemental composition of virus particles: implications for marine biogeochemical cycles. Nature Reviews Microbiology 12, 519–528.

Kanehisa, M., and Goto, S. (2000). KEGG: Kyoto Encyclopedia of Genes and Genomes. Nucleic Acids Res 28, 27–30.

Kanehisa, M., Sato, Y., and Morishima, K. (2016). BlastKOALA and GhostKOALA: KEGG Tools for Functional Characterization of Genome and Metagenome Sequences. J. Mol. Biol. 428, 726–731.

Kaplan, I.R., and Rittenberg, S.C. (1964). Microbiological Fractionation of Sulphur Isotopes. Journal of General Microbiology 34, 195–212.

Katoh, K., and Standley, D.M. (2013). MAFFT Multiple Sequence Alignment Software Version 7: Improvements in Performance and Usability. Mol Biol Evol 30, 772–780.

Kessler, D. (2006). Enzymatic activation of sulfur for incorporation into biomolecules in prokaryotes. FEMS Microbiol Rev 30, 825–840.

Kieft, K., Zhou, Z., and Anantharaman, K. (2020). VIBRANT: automated recovery, annotation and curation of microbial viruses, and evaluation of viral community function from genomic sequences. Microbiome 8, 90.

Knauer, S.H., Hartl-Spiegelhauer, O., Schwarzinger, S., Hänzelmann, P., and Dobbek, H. (2012). The Fe(II)/α-ketoglutarate-dependent taurine dioxygenases from Pseudomonas putida and Escherichia coli are tetramers. The FEBS Journal 279, 816–831.

Komoto, J., Yamada, T., Takata, Y., Markham, G.D., and Takusagawa, F. (2004). Crystal Structure of the S-Adenosylmethionine Synthetase Ternary Complex: A Novel Catalytic Mechanism of S-Adenosylmethionine Synthesis from ATP and Met,. Biochemistry 43, 1821–1831.

Kuang, Y.-S., Lu, J.-H., Li, S.-H., Li, J.-H., Yuan, M.-Y., He, J.-R., Chen, N.-N., Xiao, W.-Q., Shen, S.-Y., Qiu, L., et al. (2017). Connections between the human gut microbiome and gestational diabetes mellitus. Gigascience 6.

Kuijpers, N.G., Solis-Escalante, D., Bosman, L., van den Broek, M., Pronk, J.T., Daran, J.-M., and Daran-Lapujade, P. (2013). A versatile, efficient strategy for assembly of multi-fragment expression vectors in Saccharomyces cerevisiae using 60 bp synthetic recombination sequences. Microbial Cell Factories 12, 47.

Langmead, B., and Salzberg, S.L. (2012). Fast gapped-read alignment with Bowtie 2. Nat. Methods 9, 357–359.

Larsen, J.M. (2017). The immune response to Prevotella bacteria in chronic inflammatory disease. Immunology 151, 363–374.

Li, W., and Godzik, A. (2006). Cd-hit: a fast program for clustering and comparing large sets of protein or nucleotide sequences. Bioinformatics 22, 1658–1659.

Li, Y., Hugenholtz, J., Sybesma, W., Abee, T., and Molenaar, D. (2005). Using Lactococcus lactis for glutathione overproduction. Appl. Microbiol. Biotechnol. 67, 83–90.

Lillehaug, D. (1997). An improved plaque assay for poor plaque-producing temperate lactococcal bacteriophages. Journal of Applied Microbiology 83, 85–90.

Linz, A.M., He, S., Stevens, S.L.R., Anantharaman, K., Rohwer, R.R., Malmstrom, R.R., Bertilsson, S., and McMahon, K.D. (2018). Freshwater carbon and nutrient cycles revealed through reconstructed population genomes. PeerJ 6, e6075.

Linz, A.M., Aylward, F.O., Bertilsson, S., and McMahon, K.D. (2020). Time-series metatranscriptomes reveal conserved patterns between phototrophic and heterotrophic microbes in diverse freshwater systems. Limnology and Oceanography 65, S101–S112.

Lira, N.P.V. de, Pauletti, B.A., Marques, A.C., Perez, C.A., Caserta, R., Souza, A.A. de, Vercesi, A.E., Leme, A.F.P., and Benedetti, C.E. (2018). BigR is a sulfide sensor that regulates a sulfur transferase/dioxygenase required for aerobic respiration of plant bacteria under sulfide stress. Scientific Reports 8, 3508.

Liu, X., Jiang, H., Gu, Z., and Roberts, J.W. (2013). High-resolution view of bacteriophage lambda gene expression by ribosome profiling. Proc Natl Acad Sci U S A 110, 11928–11933.

Livak, K.J., and Schmittgen, T.D. (2001). Analysis of relative gene expression data using realtime quantitative PCR and the 2(-Delta Delta C(T)) Method. Methods 25, 402–408.

Lucke, K. (2006). Prevalence of Bacteroides and Prevotella spp. in ulcerative colitis. Journal of Medical Microbiology 55, 617–624.

Ma, H., Cheng, X., Li, G., Chen, S., Quan, Z., Zhao, S., and Niu, L. (2000). The influence of hydrogen sulfide on corrosion of iron under different conditions. Corrosion Science 42, 1669–1683.

Mahmoudabadi, G., Milo, R., and Phillips, R. (2017). Energetic cost of building a virus. PNAS 114, E4324–E4333.

Mann, N.H., Cook, A., Millard, A., Bailey, S., and Clokie, M. (2003). Bacterial photosynthesis genes in a virus. Nature 424, 741.

Manojlović, L.M. (2015). Photometry-based estimation of the total number of stars in the Universe. Applied Optics 54, 6589.

Marchler-Bauer, A., Bo, Y., Han, L., He, J., Lanczycki, C.J., Lu, S., Chitsaz, F., Derbyshire, M.K., Geer, R.C., Gonzales, N.R., et al. (2017). CDD/SPARCLE: functional classification of proteins via subfamily domain architectures. Nucleic Acids Res 45, D200–D203.

Markowitz, V.M., Chen, I.-M.A., Chu, K., Szeto, E., Palaniappan, K., Pillay, M., Ratner, A., Huang, J., Pagani, I., Tringe, S., et al. (2014). IMG/M 4 version of the integrated metagenome comparative analysis system. Nucleic Acids Res 42, D568–D573.

Martín, R., Miquel, S., Benevides, L., Bridonneau, C., Robert, V., Hudault, S., Chain, F., Berteau, O., Azevedo, V., Chatel, J.M., et al. (2017). Functional Characterization of Novel Faecalibacterium prausnitzii Strains Isolated from Healthy Volunteers: A Step Forward in the Use of F. prausnitzii as a Next-Generation Probiotic. Front Microbiol 8.

Moran, M.A., and Durham, B.P. (2019). Sulfur metabolites in the pelagic ocean. Nature Reviews Microbiology 17, 665–678.

Morra, M.J., and Dick, W.A. (1991). Mechanisms of H2S Production from Cysteine and Cystine by Microorganisms Isolated from Soil by Selective Enrichment. Appl Environ Microbiol 57, 1413–1417.

Nambi, S., Long, J.E., Mishra, B.B., Baker, R., Murphy, K.C., Olive, A.J., Nguyen, H.P., Shaffer, S.A., and Sassetti, C.M. (2015). The oxidative stress network of Mycobacterium tuberculosis reveals coordination between radical detoxification systems. Cell Host Microbe 17, 829–837.

Nayfach, S., Shi, Z.J., Seshadri, R., Pollard, K.S., and Kyrpides, N.C. (2019). New insights from uncultivated genomes of the global human gut microbiome. Nature 1.

Norman, J.M., Handley, S.A., Baldridge, M.T., Droit, L., Liu, C.Y., Keller, B.C., Kambal, A., Monaco, C.L., Zhao, G., Fleshner, P., et al. (2015). Disease-Specific Alterations in the Enteric Virome in Inflammatory Bowel Disease. Cell 160, 447–460.

O’Leary, N.A., Wright, M.W., Brister, J.R., Ciufo, S., Haddad, D., McVeigh, R., Rajput, B., Robbertse, B., Smith-White, B., Ako-Adjei, D., et al. (2016). Reference sequence (RefSeq) database at NCBI: current status, taxonomic expansion, and functional annotation. Nucleic Acids Res 44, D733–D745.

Olm, M.R., Brown, C.T., Brooks, B., Firek, B., Baker, R., Burstein, D., Soenjoyo, K., Thomas, B.C., Morowitz, M., and Banfield, J.F. (2017a). Identical bacterial populations colonize premature infant gut, skin, and oral microbiomes and exhibit different in situ growth rates. Genome Res.

Olm, M.R., Brown, C.T., Brooks, B., and Banfield, J.F. (2017b). dRep: a tool for fast and accurate genomic comparisons that enables improved genome recovery from metagenomes through de-replication. The ISME Journal 11, 2864–2868.

Paez-Espino, D., Eloe-Fadrosh, E.A., Pavlopoulos, G.A., Thomas, A.D., Huntemann, M., Mikhailova, N., Rubin, E., Ivanova, N.N., and Kyrpides, N.C. (2016). Uncovering Earth’s virome. Nature 536, 425–430.

Paez-Espino, D., Chen, I.-M.A., Palaniappan, K., Ratner, A., Chu, K., Szeto, E., Pillay, M., Huang, J., Markowitz, V.M., Nielsen, T., et al. (2017). IMG/VR: a database of cultured and uncultured DNA Viruses and retroviruses. Nucleic Acids Res. 45, D457–D465.

Pal, V.K., Bandyopadhyay, P., and Singh, A. (2018). Hydrogen Sulfide in Physiology and Pathogenesis of Bacteria and Viruses. IUBMB Life 70, 393–410.

Park, S., and Imlay, J.A. (2003). High Levels of Intracellular Cysteine Promote Oxidative DNA Damage by Driving the Fenton Reaction. J. Bacteriol. 185, 1942–1950.

Pasolli, E., Asnicar, F., Manara, S., Zolfo, M., Karcher, N., Armanini, F., Beghini, F., Manghi, P., Tett, A., Ghensi, P., et al. (2019). Extensive Unexplored Human Microbiome Diversity Revealed by Over 150,000 Genomes from Metagenomes Spanning Age, Geography, and Lifestyle. Cell 176, 649–662.e20.

Patrascu, O., Béguet-Crespel, F., Marinelli, L., Le Chatelier, E., Abraham, A.-L., Leclerc, M., Klopp, C., Terrapon, N., Henrissat, B., Blottière, H.M., et al. (2017). A fibrolytic potential in the human ileum mucosal microbiota revealed by functional metagenomic. Sci Rep 7.

Peng, H., Shen, J., Edmonds, K.A., Luebke, J.L., Hickey, A.K., Palmer, L.D., Chang, F.-M.J., Bruce, K.A., Kehl-Fie, T.E., Skaar, E.P., et al. (2017). Sulfide Homeostasis and Nitroxyl Intersect via Formation of Reactive Sulfur Species in Staphylococcus aureus. MSphere 2.

Pfleiderer, A., Mishra, A.K., Lagier, J.-C., Robert, C., Caputo, A., Raoult, D., and Fournier, P.-E. (2014). Non-contiguous finished genome sequence and description of Alistipes ihumii sp. nov. Standards in Genomic Sciences 9, 1221.

Poyraz, Ö., Brunner, K., Lohkamp, B., Axelsson, H., Hammarström, L.G.J., Schnell, R., and Schneider, G. (2015). Crystal Structures of the Kinase Domain of the Sulfate-Activating Complex in Mycobacterium tuberculosis. PLoS One 10.

Propst-Ricciuti, C. (1976). The Effect of Host-Cell Starvation on Virus-induced Lysis by MS2 Bacteriophage. Journal of General Virology 31, 323–330.

Qin, J., Li, R., Raes, J., Arumugam, M., Burgdorf, K.S., Manichanh, C., Nielsen, T., Pons, N., Levenez, F., Yamada, T., et al. (2010). A human gut microbial gene catalog established by metagenomic sequencing. Nature 464, 59–65.

Rahlff, J., Turzynski, V., Esser, S.P., Monsees, I., Bornemann, T.L.V., Figueroa-Gonzalez, P.A., Schulz, F., Woyke, T., Klingl, A., Moraru, C., et al. (2020). Genome-informed microscopy reveals infections of uncultivated carbon-fixing archaea by lytic viruses in Earth’s crust. BioRxiv 2020.07.22.215848.

Rambaut, A. (2009). FigTree version 1.4.3.

Roux, S., Hawley, A.K., Beltran, M.T., Scofield, M., Schwientek, P., Stepanauskas, R., Woyke, T., Hallam, S.J., and Sullivan, M.B. (2014). Ecology and evolution of viruses infecting uncultivated SUP05 bacteria as revealed by single-cell- and meta-genomics. ELife Sciences 3, e03125.

Roux, S., Enault, F., Hurwitz, B.L., and Sullivan, M.B. (2015). VirSorter: mining viral signal from microbial genomic data. PeerJ 3.

Roux, S., Brum, J.R., Dutilh, B.E., Sunagawa, S., Duhaime, M.B., Loy, A., Poulos, B.T., Solonenko, N., Lara, E., Poulain, J., et al. (2016). Ecogenomics and potential biogeochemical impacts of globally abundant ocean viruses. Nature 537, 689–693.

Schirmer, M., Franzosa, E.A., Lloyd-Price, J., McIver, L.J., Schwager, R., Poon, T.W., Ananthakrishnan, A.N., Andrews, E., Barron, G., Lake, K., et al. (2018). Dynamics of metatranscription in the inflammatory bowel disease gut microbiome. Nature Microbiology 3, 337–346.

Schulz, F., Andreani, J., Francis, R., Boudjemaa, H., Khalil, J.Y.B., Lee, J., Scola, B.L., and Woyke, T. (2020). Advantages and Limits of Metagenomic Assembly and Binning of a Giant Virus. MSystems 5.

Seemann, T. (2014). Prokka: rapid prokaryotic genome annotation. Bioinformatics 30, 2068–2069.

Sernova, N.V., and Gelfand, M.S. (2008). Identification of replication origins in prokaryotic genomes. Brief Bioinform 9, 376–391.

Shimizu, T., Shen, J., Fang, M., Zhang, Y., Hori, K., Trinidad, J.C., Bauer, C.E., Giedroc, D.P., and Masuda, S. (2017). Sulfide-responsive transcriptional repressor SqrR functions as a master regulator of sulfide-dependent photosynthesis. Proceedings of the National Academy of Sciences 114, 2355–2360.

Sim, M.S., Ogata, H., Lubitz, W., Adkins, J.F., Sessions, A.L., Orphan, V.J., and McGlynn, S.E. (2019). Role of APS reductase in biogeochemical sulfur isotope fractionation. Nature Communications 10, 44.

Stamatakis, A. (2014). RAxML version 8: a tool for phylogenetic analysis and post-analysis of large phylogenies. Bioinformatics 30, 1312–1313.

Sullivan, M.B., Lindell, D., Lee, J.A., Thompson, L.R., Bielawski, J.P., and Chisholm, S.W. (2006). Prevalence and Evolution of Core Photosystem II Genes in Marine Cyanobacterial Viruses and Their Hosts. PLoS Biology 4, e234.

Sullivan, M.J., Petty, N.K., and Beatson, S.A. (2011). Easyfig: a genome comparison visualizer. Bioinformatics 27, 1009–1010.

Suttle, C.A. (2005). Viruses in the sea. Nature 437, 356–361.

Suttle, C.A. (2007). Marine viruses — major players in the global ecosystem. Nature Reviews Microbiology 5, 801–812.

Tam, W., Pell, L.G., Bona, D., Tsai, A., Dai, X.X., Edwards, A.M., Hendrix, R.W., Maxwell, K.L., and Davidson, A.R. (2013). Tail tip proteins related to bacteriophage λ gpL coordinate an iron-sulfur cluster. J. Mol. Biol. 425, 2450–2462.

Tatusova, T., DiCuccio, M., Badretdin, A., Chetvernin, V., Nawrocki, E.P., Zaslavsky, L., Lomsadze, A., Pruitt, K.D., Borodovsky, M., and Ostell, J. (2016). NCBI prokaryotic genome annotation pipeline. Nucleic Acids Res 44, 6614–6624.

Thode, H.G., Macnamara, J., and Fleming, W.H. (1953). Sulphur isotope fractionation in nature and geological and biological time scales. Geochimica et Cosmochimica Acta 3, 235–243.

Thompson, L.R., Zeng, Q., Kelly, L., Huang, K.H., Singer, A.U., Stubbe, J., and Chisholm, S.W. (2011). Phage auxiliary metabolic genes and the redirection of cyanobacterial host carbon metabolism. PNAS 108, E757–E764.

Trubl, G., Jang, H.B., Roux, S., Emerson, J.B., Solonenko, N., Vik, D.R., Solden, L., Ellenbogen, J., Runyon, A.T., Bolduc, B., et al. (2018). Soil Viruses Are Underexplored Players in Ecosystem Carbon Processing. MSystems 3, e00076–18.

UniProt Consortium, T. (2018). UniProt: the universal protein knowledgebase. Nucleic Acids Res 46, 2699–2699.

Veiga, P., Pons, N., Agrawal, A., Oozeer, R., Guyonnet, D., Brazeilles, R., Faurie, J.-M., van Hylckama Vlieg, J.E.T., Houghton, L.A., Whorwell, P.J., et al. (2014). Changes of the human gut microbiome induced by a fermented milk product. Sci Rep 4.

Villion, M., Chopin, M.-C., Deveau, H., Ehrlich, S.D., Moineau, S., and Chopin, A. (2009). P087, a lactococcal phage with a morphogenesis module similar to an Enterococcus faecalis prophage. Virology 388, 49–56.

Voordouw, G., Armstrong, S.M., Reimer, M.F., Fouts, B., Telang, A.J., Shen, Y., and Gevertz, D. (1996). Characterization of 16S rRNA genes from oil field microbial communities indicates the presence of a variety of sulfate-reducing, fermentative, and sulfide-oxidizing bacteria. Appl Environ Microbiol 62, 1623–1629.

Wacey, D., Kilburn, M.R., Saunders, M., Cliff, J., and Brasier, M.D. (2011). Microfossils of sulphur-metabolizing cells in 3.4-billion-year-old rocks of Western Australia. Nature Geoscience 4, 698–702.

Wilhelm, S.W., and Suttle, C.A. (1999). Viruses and Nutrient Cycles in the Sea. BioScience 49, 8.

Xia, Y., Lü, C., Hou, N., Xin, Y., Liu, J., Liu, H., and Xun, L. (2017). Sulfide production and oxidation by heterotrophic bacteria under aerobic conditions. The ISME Journal 11, 2754–2766.

Yang, Y., Xu, G., Liang, J., He, Y., Xiong, L., Li, H., Bartlett, D., Deng, Z., Wang, Z., and Xiao, X. (2017). DNA Backbone Sulfur-Modification Expands Microbial Growth Range under Multiple Stresses by its anti-oxidation function. Scientific Reports 7, 3516.

Yeeles, J.T.P., Cammack, R., and Dillingham, M.S. (2009). An Iron-Sulfur Cluster Is Essential for the Binding of Broken DNA by AddAB-type Helicase-Nucleases. J Biol Chem 284, 7746–7755.

Yin, J., Ren, W., Yang, G., Duan, J., Huang, X., Fang, R., Li, C., Li, T., Yin, Y., Hou, Y., et al. (2016). l-Cysteine metabolism and its nutritional implications. Molecular Nutrition & Food Research 60, 134–146.

