## Supplemental Figures S1-S9 for "Virus-associated organosulfur metabolism in human and environmental systems"

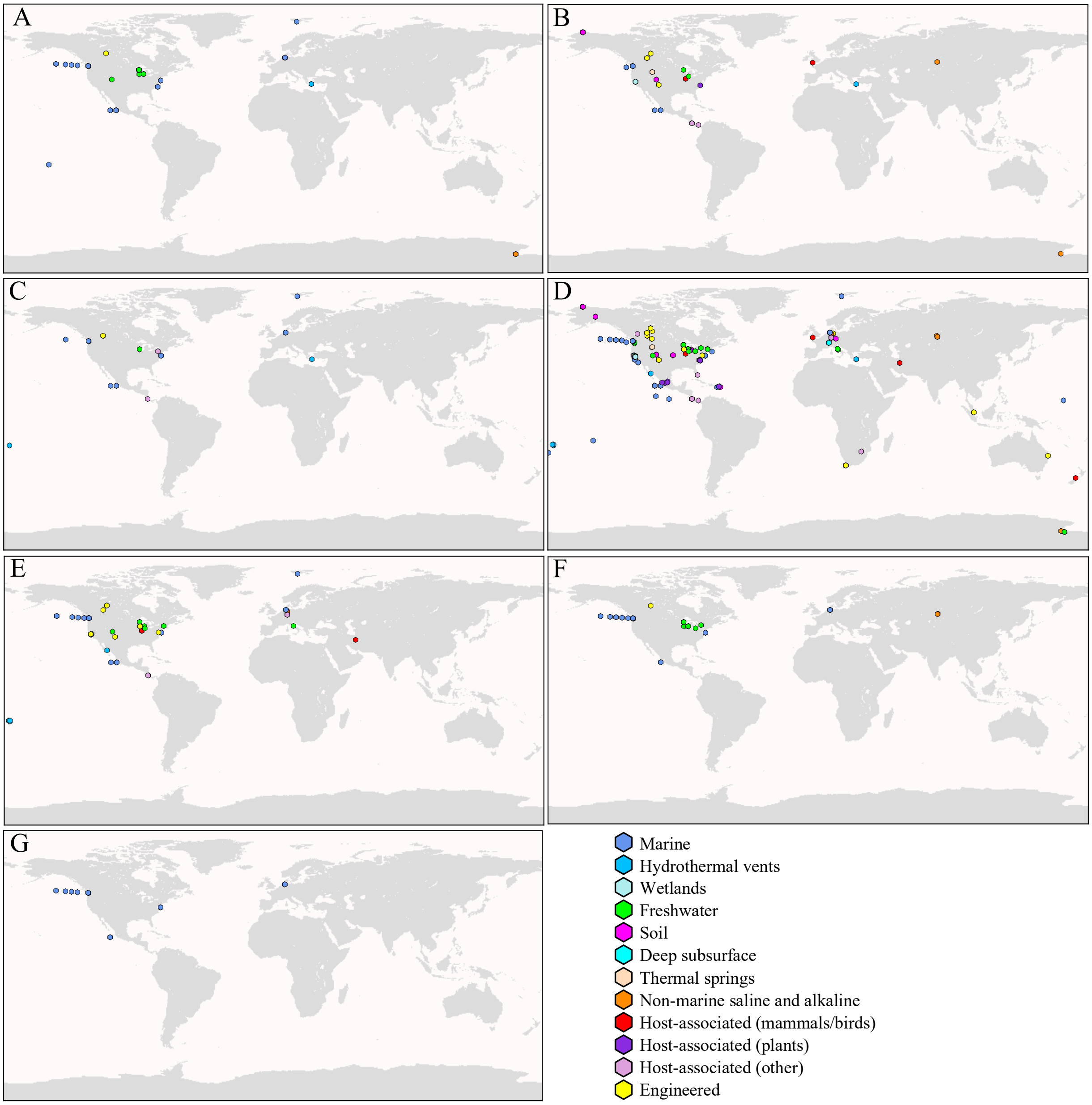


**Figure S1. Distribution of individual AMGs in the environment, related to Figure 2.** Global distribution of viral populations encoding (A) *cysC*, (B) *cysH*, (C) *cysK*, (D) *dcm*, (E) *metK*, (F) *tauD* or (G) *msmA* identified on the IMG/VR database, color coordinated by environment classification.

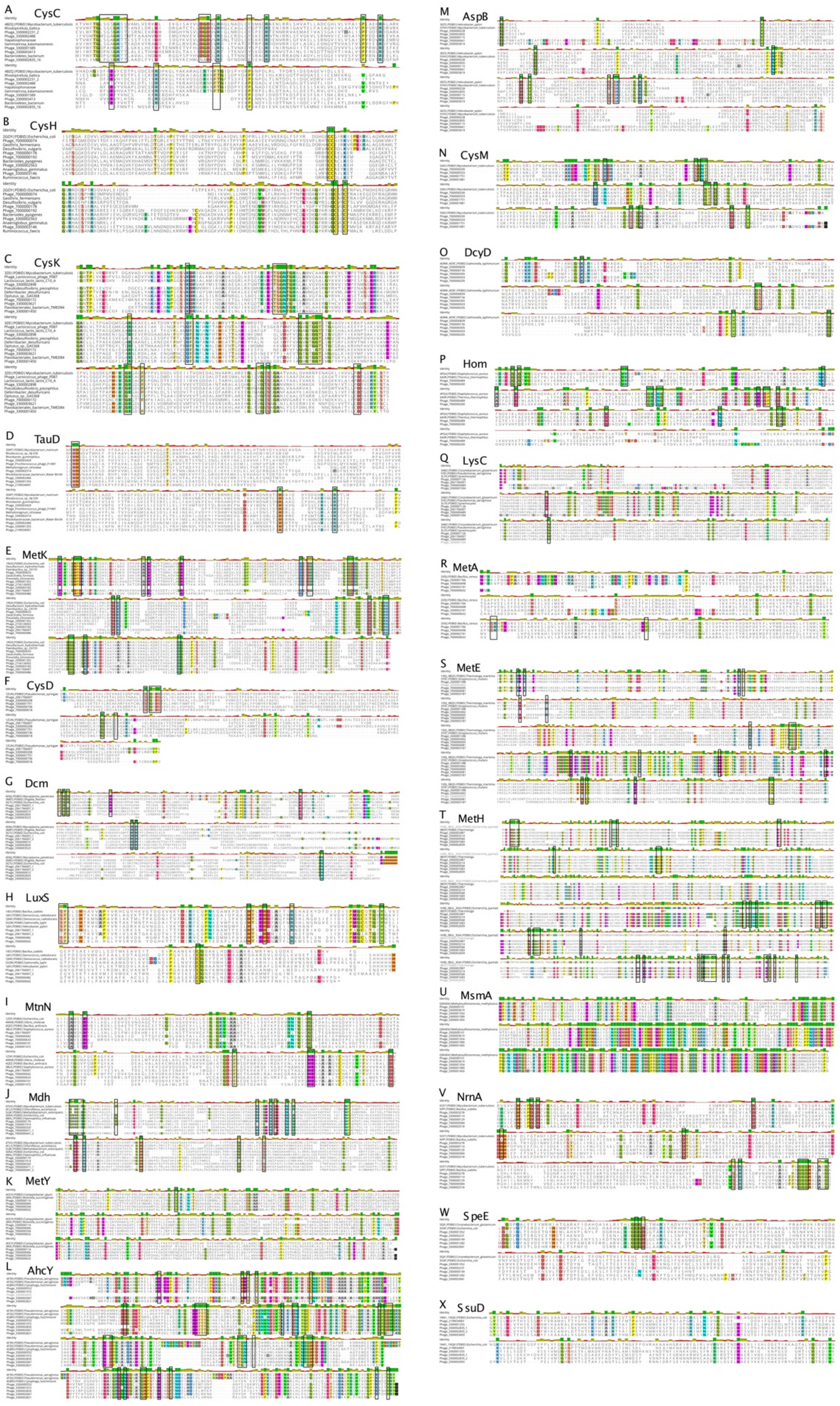
A CysC
B CysH
C CysK
D TauD
E MetK
F CysD
G Dcm
H LuxS
I MtnN
J Mdh
K MetY
L AhcY
M AspB
N CysM
O DcyD
P Hom
Q LysC
R MetA
S MetE
T MetH
U MsmA
V NrnA
W SpeE
X SsuD

**Figure S2. Amino acid alignments of proteins encoded by AMGs, related to Figure 1 and Table 1.** Alignment of representative viral and bacterial sequences for all AMGs with abundances greater than five. Viruses are indicated by the preface "Phage" followed by their respective IMG Taxon Object ID number. See Table S1 for full genome names. Refer to Figurer S4 for phylogeny of the represented sequences for A-E. Highlighted amino acids indicate conservation in >85% of sequences shown. Black boxes indicate amino acid residues that are biochemically verified as functional according to the given PDB reference sequence.

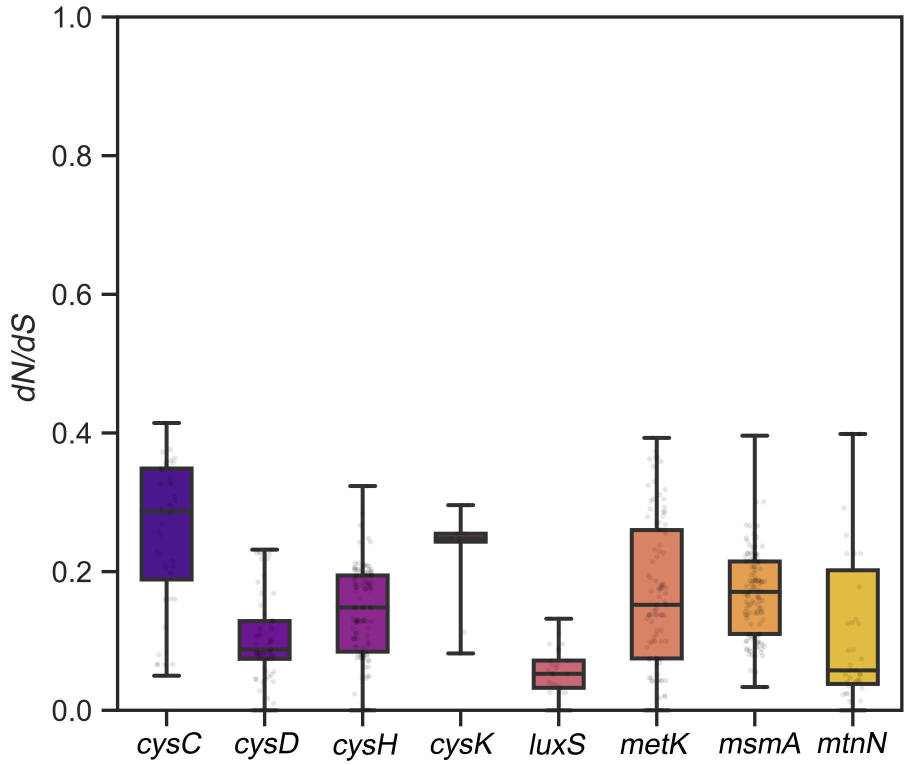


**Figure S3. *dN/dS* ratio calculations for cysK, cysC, cysD, cysH, tauD, msmA, metK, mtmN and luxS AMG pairs, related to Figures 1, 4 and 5.** Each data point overlaid on the standard box plot represents a single AMG pair. The figure was generated using seaborn and matplotlib.

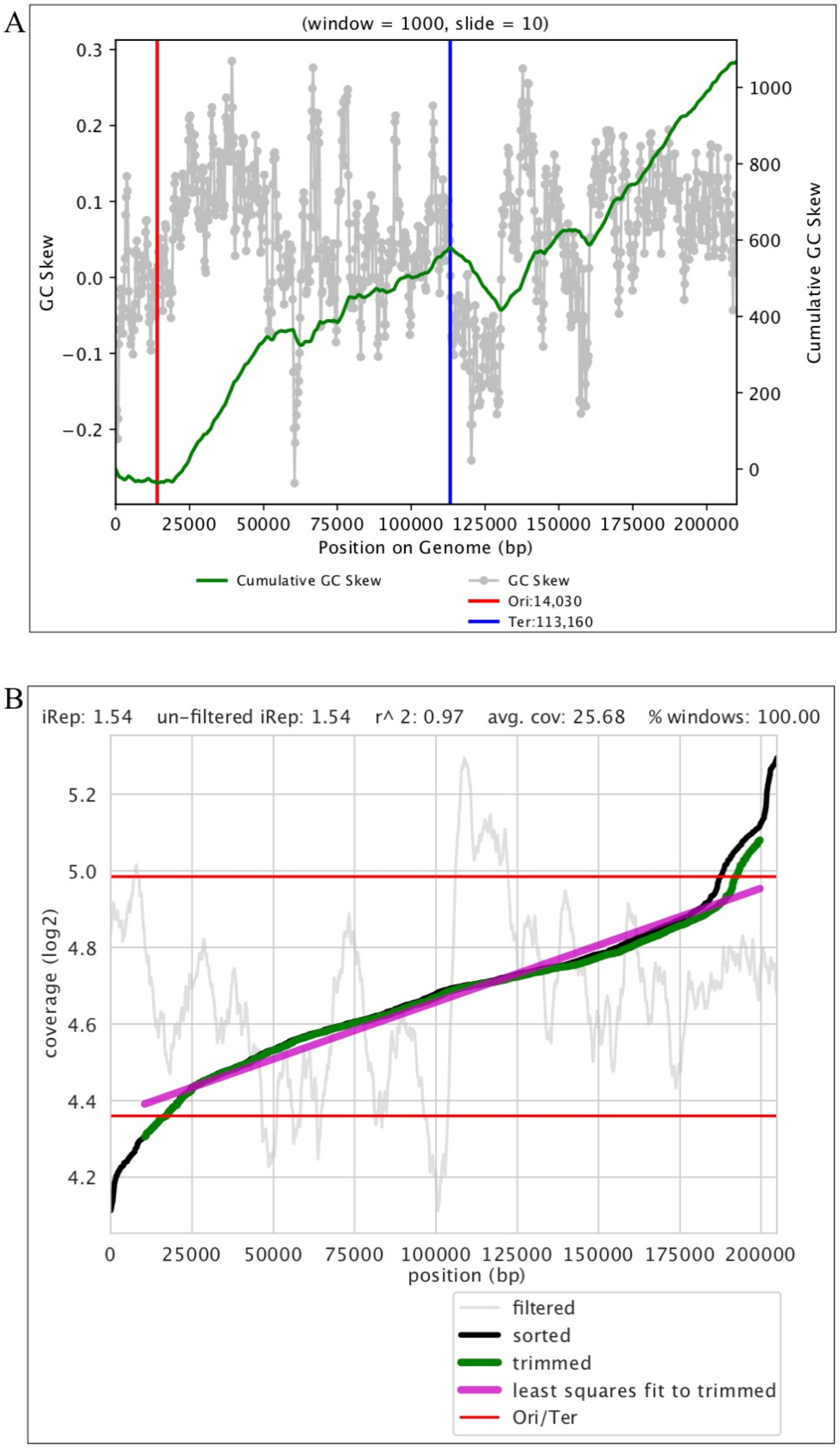


**Figure S4. Genome and growth statistics of a complete virus identified to express *cysC* in Lake Mendota, WI, related to Figure 6.** The (A) GC-skew and (B) index of replication for a complete, circular virus identified in Lake Mendota, WI. GC-skew and replication statistics indicate that the virus was actively replicating at time of sampling and likely undergoes rolling circle replication.

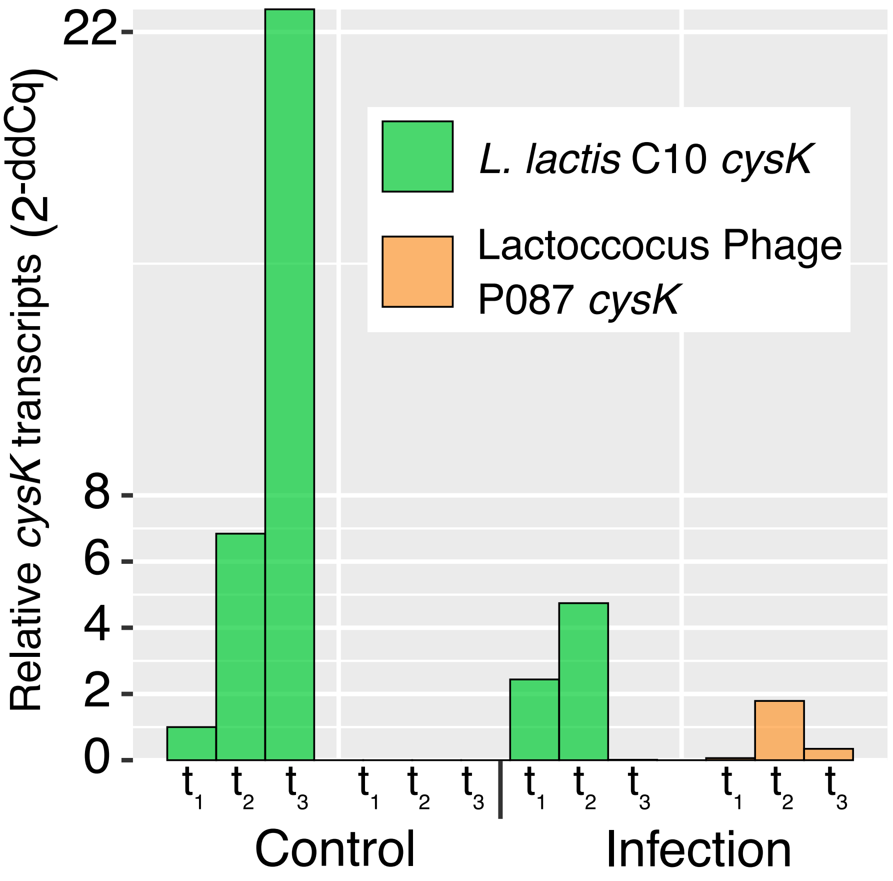


**Figure S5. qPCR results of *cysK* transcript relative abundance for *L. lactis* C10 and phage P087, related to Figure 3.** Relative transcript abundance is provided in the normalized $2^{-ddCq}$ metric. Control is *L. lactis* C10 (host) alone and infection includes host plus phage P087. The times shown are $t_1$ (15 minutes), $t_2$ (60 minutes) and $t_3$ (120 minutes).

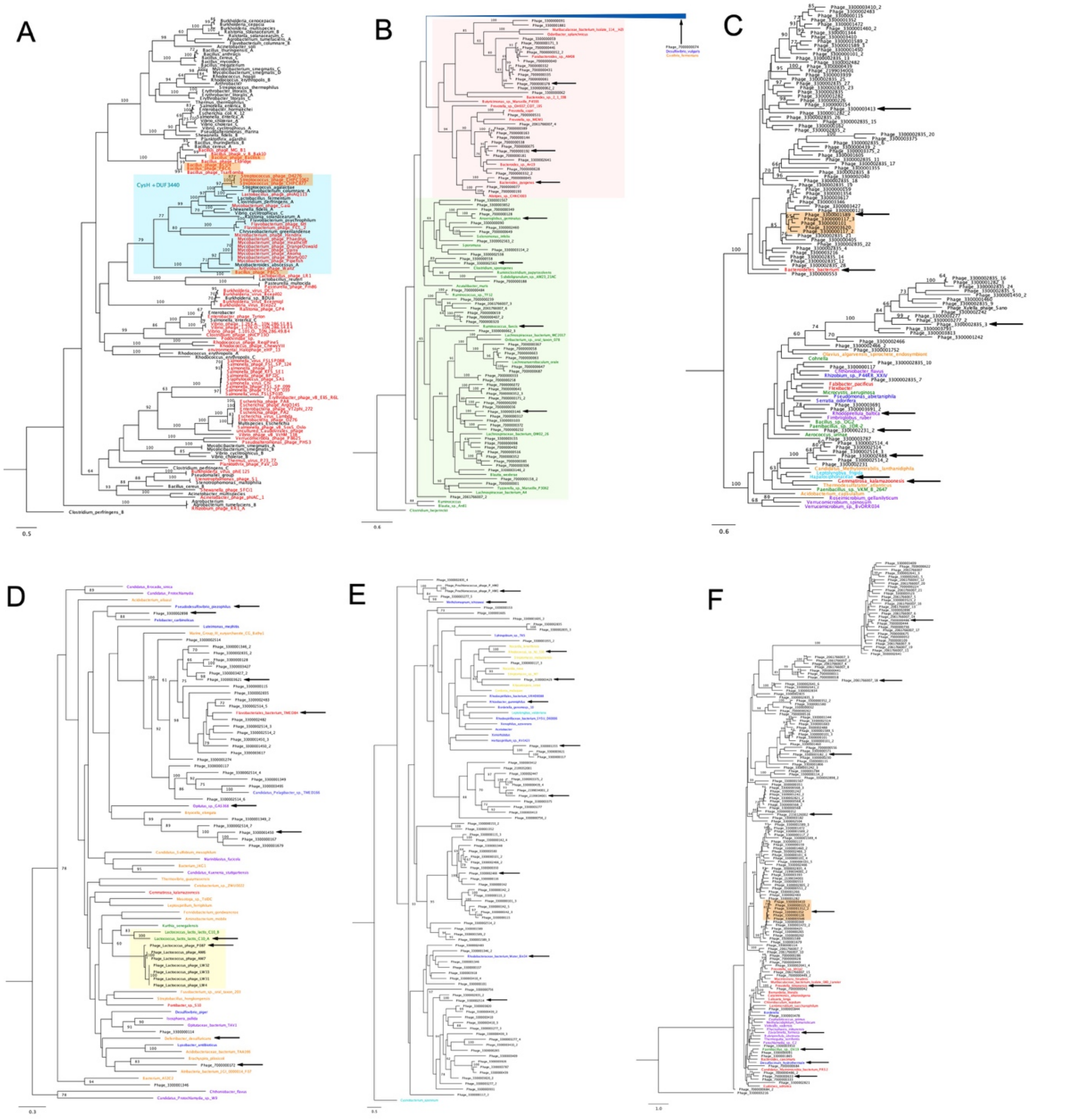


**Figure S6. Phylogeny of viral AMG encoded protein sequences, related to Figures 1, 4 and 5**. (A) Phylogenetic tree for CysH of complete viruses with known bacterial hosts. Viruses are in red and bacteria are in black, and proteins with an additional DUF4440 domain are highlighted in blue. Bacteria with multiple copies of *cysH* are appended with a letter ("A" through "D"). Refer to Table S1 for virus-host associations. Also shown are phylogenetic trees of uncultivated viruses with bacterial homologs and select cultivated viruses for (B) CysH (red and green highlighting refers to putative virus-host associations; compressed blue clade contains 36 viruses and 89 bacteria from several phyla), (C) CysC, (D) CysK (yellow highlighting refers to known virus-host associations), (E) TauD and (F) MetK. For trees (B-F) colored names refer to viruses (black), Bacteroidetes and other members of the FCB superphylum (red), Cyanobacteria (cyan), Verrucomicrobia and Planctomycetes and other members of the PVC superphylum (purple), Actinobacteria (yellow), and all other phyla in orange. For all trees, bootstrap values greater than 60 are shown, orange highlighting indicates respective genomes used for comparative genomics (see Figures 4 and S6), and arrows indicate sequences used for protein alignments (see Figure S2).

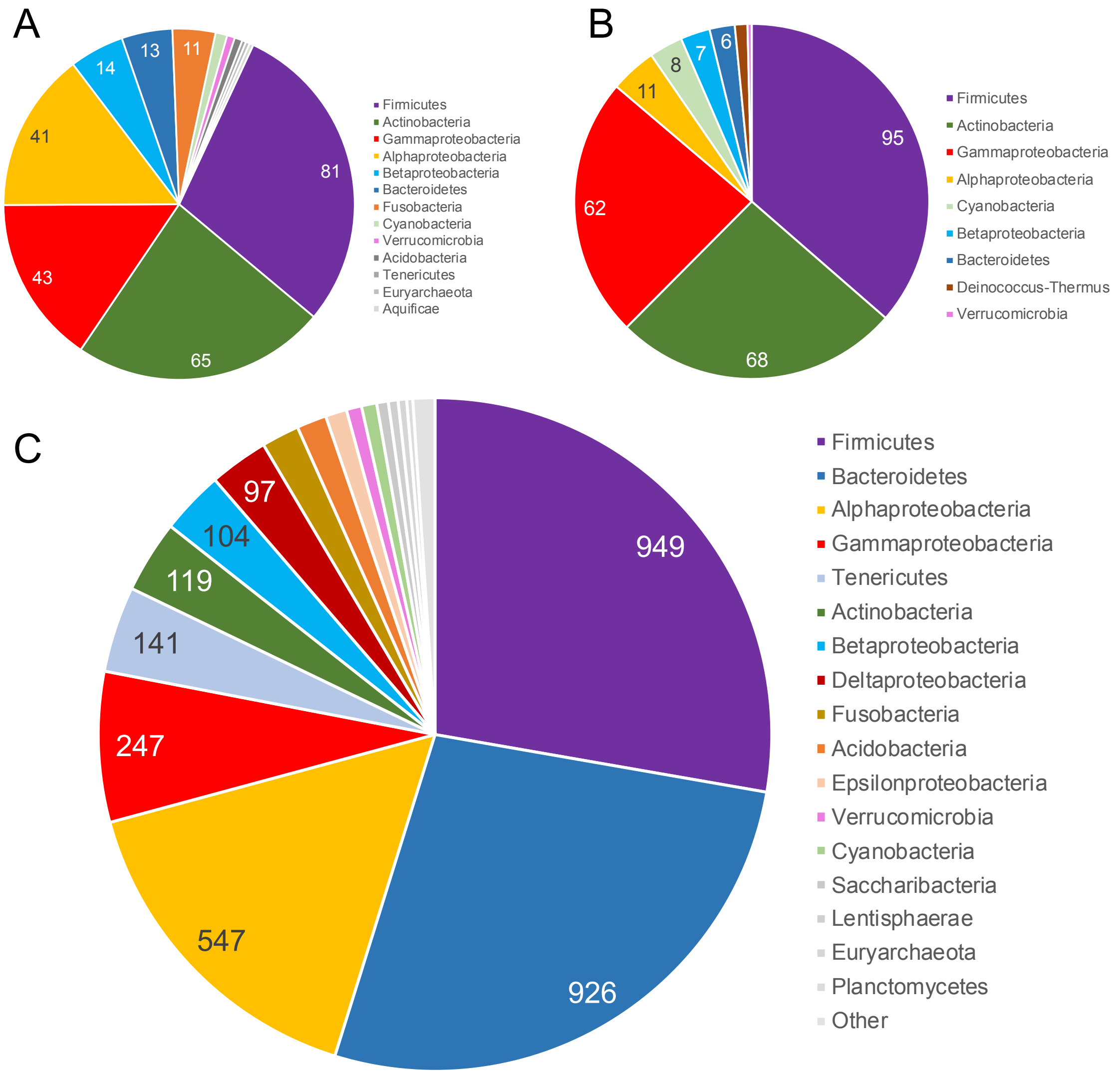


**Figure S7. Taxonomic classification of AMGs, related to Figure 2**. Inferred taxonomic classification at the phylum level of AMG-encoded protein sequences on NCBI-derived (A) viruses and (B) taxonomy of their known hosts, showing similar proportionality. (C) Inferred taxonomic classification at the phylum level of AMG-encoded protein sequences on IMG/VR-derived viruses.

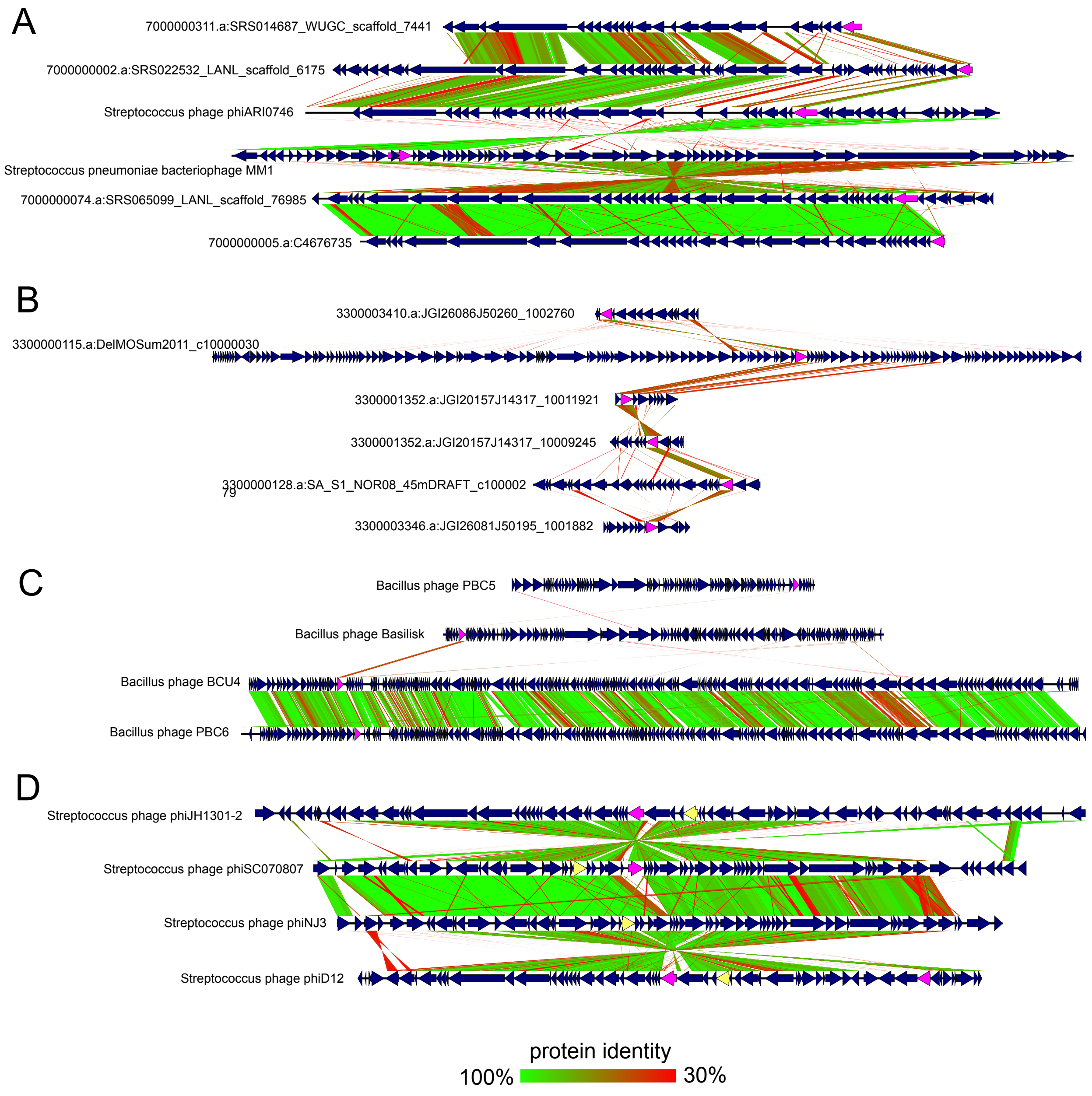


**Figure S8. Genome comparisons of viruses encoding AMGs, related to Figure 4**. Comparisons of (A) incomplete viruses and complete *Streptococcus pneumoniae* viruses encoding *dcm* (pink), (B) incomplete viruses encoding *metK* (pink), (C) complete *Bacillus cereus* viruses encoding *cysH* (pink), and (D) complete *Streptococcus suis* viruses encoding *metK* (yellow) and *dcm* (pink). For all comparisons, predicted open readings frames are annotated by dark blue arrows and genomes are connected with lines according to tblastx identity. Refer to Figure S6 for phylogeny of AMGs for (B) and (C) (orange highlighting).

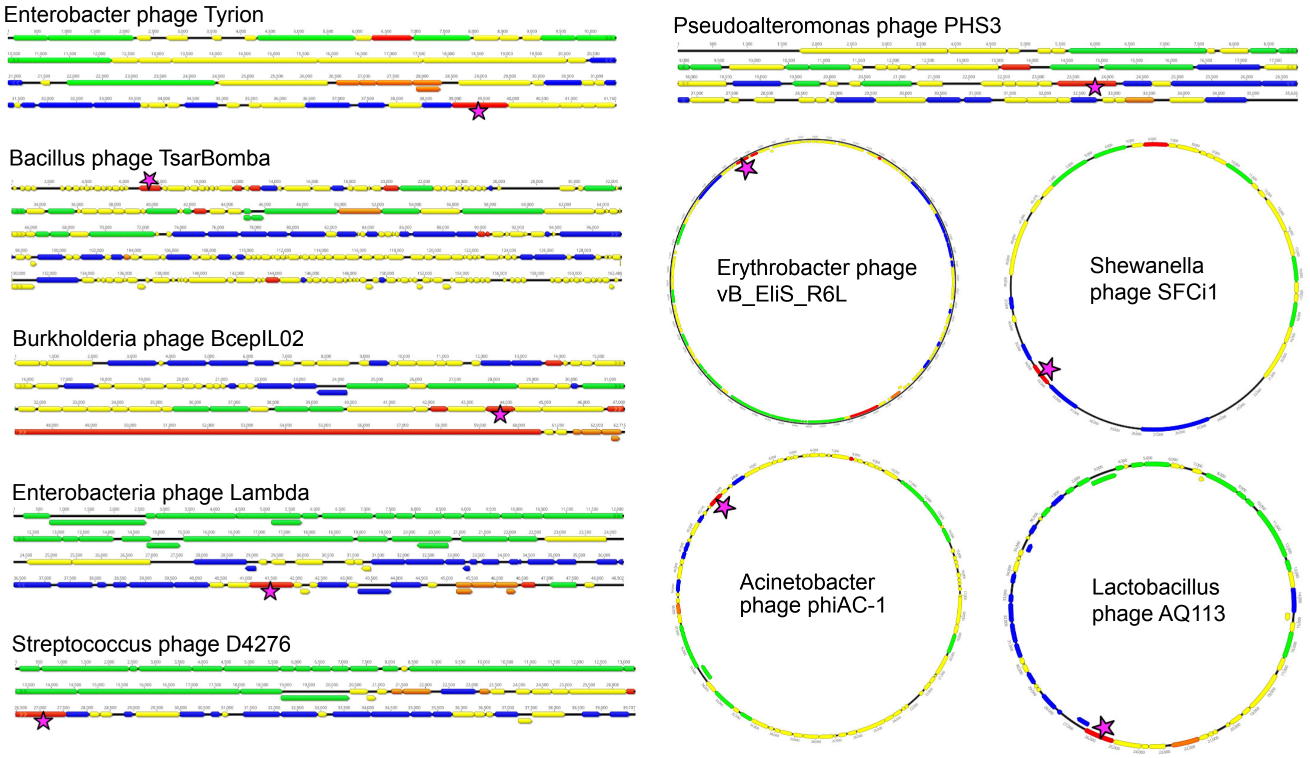


**Figure S9. Genome organization of *cysH*-encoding viruses, related to Figure 5.** Organization of linear and circular complete viral genomes that encode *cysH*. Arrows indicate open reading frames and are annotated by general function: virion structural assembly (green), auxiliary metabolism and general functions (red), nucleotide metabolism and genome replication (blue), lysis (orange) and unknown function (yellow). Pink stars indicate the location of *cysH*. Refer to Table S1 for virus details.
